# Epigenetic cell memory: The gene’s inner chromatin modification circuit

**DOI:** 10.1101/2022.02.02.476953

**Authors:** Simone Bruno, Ruth J. Williams, Domitilla Del Vecchio

## Abstract

Epigenetic cell memory allows distinct gene expression patterns to persist in different cell types despite a common genotype. Although different patterns can be maintained by the concerted action of transcription factors (TFs), it was proposed that long-term persistence hinges on chromatin state. Here, we study how the dynamics of chromatin state affect memory, and focus on a biologically motivated circuit motif, among histones and DNA modifications, that mediates the action of TFs on gene expression. Memory arises from time-scale separation among three circuit’s constituent processes: basal erasure, auto and cross-catalysis, and recruited erasure of modifications. When the two latter processes are sufficiently faster than the former, the circuit exhibits bistability and hysteresis, allowing active and repressed gene states to coexist and persist after TF stimulus removal. The duration of memory is stochastic with a mean value that increases as time-scale separation increases, but more so for the repressed state. This asymmetry stems from the cross-catalysis between repressive histone modifications and DNA methylation and is enhanced by the relatively slower decay rate of the latter. Nevertheless, TF-mediated positive autoregulation can rebalance this asymmetry and even confers robustness of active states to repressive stimuli. More generally, by wiring positively autoregulated chromatin modification circuits under time scale separation, long-term distinct gene expression patterns arise, which are also robust to failure in the regulatory links.

## 1 Introduction

The maintenance of distinct cell identities without any difference in genotype is a critical property of multicellular organisms. This property is captured by the word “epigenetic”, coined by Conrad H. Waddington in 1942, to broadly indicate that different cell phenotypes can be associated with the same genotype [1]. A natural question is how multicellular organisms can ensure that distinct phenotypes are safely locked-in despite sharing the same genotype. This question has been extensively investigated in systems biology [2, 3] and has been recently revisited in light of the so-called “Epigenetic Revolution” [4]. Maintenance of distinct phenotypes requires the inheritance of different gene expression patterns through subsequent cell divisions [5]. During the natural process of cell differentiation, cells go through multiple fate decisions and select among different gene expression patterns that are mutually exclusive and are robust to both intrinsic noise and TF perturbations [6]. According to classical theory, these patterns are all stable attractors of the dynamics of gene regulation networks (GRNs) enabled by TF binding to DNA. In each of these attractors, lineage-specific genes become continuously expressed and often maintain their expression through positive autoregulation [7, 8, 9]. Therefore, the GRN-centric perspective proposes that TF-enabled regulatory networks are fully capable of maintaining epigenetic memory of cell-type-specific gene expression patterns in differentiated cells.

Following an explosive number of discoveries fueling the Epigenetic Revolution, a different and somewhat contrasting view has emerged on epigenetic cell memory [4]. A key argument that detracts to the role of TF-enabled regulatory networks in maintaining memory states is that maintenance is dictated by the binding of TFs to DNA. But, this binding is disrupted upon DNA replication and cell division. Allis specifically says “This mechanism is transient in dividing cells because progression of the DNA replication fork usually disrupts these protein-DNA interactions, which then need to be re-established in the resulting daughter cells.” ([10], Chapter 22). In [11], the authors also argue that “…feedback loops can clearly enable heritable states of altered gene expression without any need to evoke chromatin. However, it is unlikely that such feedback loops alone would enable the propagation of states throughout the length of development and in the germline of complex organisms.”

It has therefore been proposed that it is the chromatin state, i.e., the extent of DNA compaction mediated by histone and DNA modifications, that enables long-term transmission to successive cell generations ([11] and [10], Chapter 3). This view is supported by the discovery of enzymatic processes that can copy histone modifications from modified histones to unmodified nearby histones, facilitating the maintenance of these modifications after DNA replication and cell division [10, 12]. It is also reinforced by the already established fact that DNA methylation is copied from the parental DNA strand to the progeny strand immediately after DNA replication [13, 14] and by the discovery of cooperative interactions between DNA methylation and histone modifications [15]. However, histone and DNA modifications result from dynamic enzymatic processes that continuously write and erase modifications and not from a static read-and-copy process. It thus remains unclear how these dynamic processes may enable long-term maintenance of gene expression states. These considerations motivate the need for a theoretical framework that includes the dynamics of chromatin state in the regulation of gene expression by TFs and determines how these dynamics affect the temporal duration of memory. This question is even more pressing for engineering persistent synthetic genetic circuits in the chromosome of mammalian cells, in which silencing is a major obstacle to keeping proper circuit function for extended temporal durations [16, 17, 18, 19].

In this paper, we create a biologically motivated chemical reaction model of the *gene’s inner chromatin modification circuit* that mediates the effect of TF inputs on gene expression through histone modifications and DNA methylation. The circuit comprises auto-catalysis of histone modifications, competitive recruitment of erasure between activating and repressive modifications, cross-catalysis between repressive histone modifications and DNA methylation, and basal erasure.

In this model, the effect of TFs on gene expression is not determined by the binding of TFs to DNA alone, but by the recruitment of the enzymes for the *de novo* establishment of either activating or repressive modifications. We first determine how time scale separation among the circuit’s constituent processes affect memory of a TF input stimulus for both active and repressed chromatin states, by analyzing the stability and probability of chromatin states, the mean temporal duration of a chromatin state, and the variability of this duration (Section 3.1). Then, we study the relative contributions of TF-mediated regulation and chromatin dynamics to memory of active and repressed gene states in two ubiquitous GRN motifs: TF-mediated positive autoregulation and mutual inhibition (Sections 3.2-3.4). Specifically, we analyze the mean temporal duration of active and repressed chromatin states and their robustness to both repetitive disruptions of TF-enabled regulatory links and to undesired input stimuli, such as due to endogenous silencing.

### Related work

Models of chromatin-mediated gene regulation, in which histone modifications, DNA methylation, and TF-mediated regulation are combined together, remain under-represented in the literature [20]. Most existing models that include chromatin modifications into gene expression regulation are based on phenomenological rules rather than on the molecular reactions that regulate chromatin state, and focus on specific biological processes, such as iPSC reprogramming [21, 22, 23, 24] or epithelial-mesenchymal transition [25], and most of them are only suitable for computational simulations [21, 22, 23]. On the opposite side of the spectrum, highly detailed mechanistic models have appeared for histone modifications alone, in which each nucleosome within a gene is modeled and simulated as a distinct unit [26]. Different from these existing works, we explicitly address how the duration of memory of a chromatin state is modulated by the topological properties of the chromatin modification circuit, by relevant biochemical parameters, and by the interplay between TF-based regulation and chromatin dynamics.

## 2 The gene has an inner chromatin modification circuit

In this paper, we focus on a circuit motif that captures the interactions among H3K4 methylation/acetylation, H3K9 methylation, and DNA methylation, and has the nucleosome as the basic unit that can carry these modifications (Figure 1(a)). H3K4 methylation (H3K4me3) typically co-exists with acetylation and is associated with active chromatin state ([10], Chapter 3 and [27]), while H3K9 methylation (H3K9me2/3) and DNA methylation are a hallmark of repressed chromatin state and, as such, are found at pluripotency genes and at lineage specific genes in terminally differentiated cells [5]. Specifically, H3K4 and H3K9 methylation are typically mutually exclusive since the enzymes that methylate H3K4 do not do so if the neighboring K9 residues are already methylated [28] and *viceversa* [29]. We therefore assume that each nucleosome carries either H3K9 methylation or H3K4 methylation but not both. We also assume that H3K4 methylation and acetylation co-exist and we represent them by an activating “A” nucleosome modification (Figure 1(a)). Furthermore, for H3K9 methylation, we lump together the two methylation states (me2 and me3) for simplicity and call them “me3” because both of them are associated with gene repression. Also, although nucleosomes carry two copies of each histone, we consider them as one unit that can carry only one histone modification.

**Figure 1:**
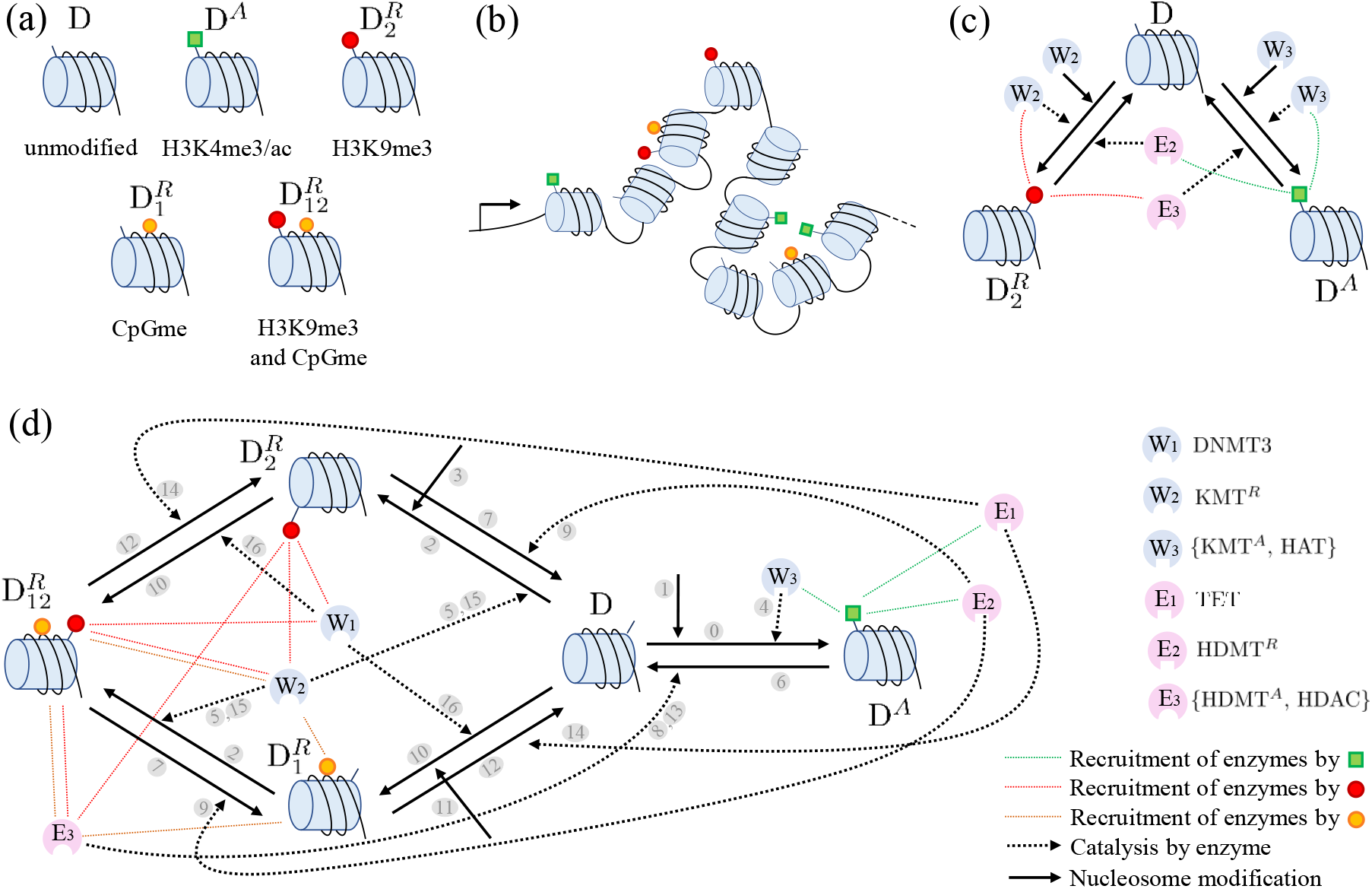
The gene’s inner chromatin modification circuit. (a) Nucleosome modifications: D (unmodified nucleosome), D^A^ (nucleosome with a activating histone modification, H3K4me3/ac), 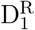 (nucleosome with only DNA methylation, CpGme), 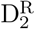 (nucleosome with only a repressive histone modification, H3K9me3) and 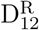 (nucleosome with both H3K9me3 and CpGme). (b) Pictorial representation of a gene with its nucleosomes carrying various modifications. The left side arrow represents the promoter. (c) Competitive interactions between opposing histone modifications (activating D^A^ and repressive 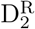), wherein each modification recruits writers of itself and erasers of the opposing modification. (d) Complete chromatin modification circuit that includes all the interactions described in Section 2 (see also SI-Figures S.2-S.3). The numbers shaded in gray on the arrows correspond to the reactions associated with the arrows, described in the main text and in Figure 2. In (c) and (d), enzymes that write (writers) and erase (erasers) each modification are represented as W_1_, W_2_, W_3_ and E_1_, E_2_, E_3_, respectively. The socket on each enzyme represents a domain that binds to protein readers of the corresponding indicated (by the dashed lines) modification, enabling the process by which each modification recruits writers or erasers to nearby histones. To distinguish the KMTs for H3K9 and H3K4, we define the writers for H3K9 methylation (Suv39H1) as KMT^*R*^ and the writers for H3K4 methylation (SETs and MLL1/2) as KMT^*A*^. Similarly, to distinguish the HDMTs for H3K9 methylation and H3K4 methylation, we define the erasers for H3K9 methylation (JMJD2A) as HDMT^*R*^ and the erasers for H3K4 methylation (JARID) as HDMT^*A*^. We use colored dotted lines to depict the recruitment process by H3K4me3/ac (green lines), H3K9me3 (red lines), and CpGme (orange lines) and we use dotted black arrows to depict the consequent effect on writing/erasing. The solid black arrow represents the nucleosome modification.

In addition to these histone modifications, DNA itself can be methylated in correspondence to the CpG dinucleotide (CpGme) [30, 10, 5]. DNA methylation is correlated with the absence of H3K4 methylation ([10], Chapter 6), since the enzymes that methylate H3K4 recognize unmethylated DNA binding motifs ([10], Chapter 1) and *viceversa* ([10], Chapter 6, and [31]). Therefore, we assume that each nucleosome can carry either one or the other but not both (Figure 1(a)). By contrast, DNA methylation can co-exist with H3K9 methylation ([10], Chapters 6, 22) and hence we allow a nucleosome to carry both modifications (Figure 1(a)). As a result, each gene will include a number of nucleosomes with different modifications (Figure 1(b)).

In the next section, we introduce the molecular mechanisms and corresponding reactions by which these modifications can be written, erased, and copied to nearby nucleosomes, which will form the chromatin modification circuit (Figure 1(c)-(d)).

### 2.1 Histone modifications

Histone modifications can be modeled by enzymatic reactions wherein an unmodified histone can be *de novo* modified by the action of writer enzymes. These enzymes are called KMTs (lysine methyltransferases) for methylation and HATs (histone acetyltransferases) for acetylation and are recruited to DNA by, among others, sequence-specific TFs ([32], Chapter 6). These modifications can be actively removed by the action of eraser enzymes. These enzymes are called HDMTs (histone demethylases) for de-methylation and HDACs (histone deacetylases) for de-acetylation. Additionally, histone modifications can be copied through a read-write process, wherein a modified histone is bound by a reader protein, which, in turn, recruits writers of the same modification [10]. We describe these in detail.

#### *De novo* establishment

As described in [10] (Chapter 21), sequence-specific transcriptional activators bind DNA and recruit HATs, such as the SAGA complex, to the promoter of a gene, which becomes acetylated. Examples of transcriptional activators that recruit HATs include Myc, GATA.1, and Gal4. The deposition of H3K4me3 then can occur co-transcriptionally as RNA Polymerase II recruits SETs, a KMT that methylates H3K4 [33] (see also [10] Chapter 3), or through the recruitment of SETs and MLL1/2 (also KMTs for H3K4) to DNA by the CxxC binding domain that specifically recognizes unmethylated DNA ([5], Chapter 7). Finally, MLLs can be recruited to specific promoters by transcriptional activators such as Oct4 [34]. H3K9me3 is established by the writer action of Suv39H1 (a KMT for H3K9), which can also be recruited to DNA by sequence-specific TFs. An example of this is the recruitment of this enzyme to GATA.1 targets by the PU.1 TF, as a means to silence GATA.1 targets and promote the myeloid lineage [35]. These modifications can be captured by the one-step enzymatic reactions ⓪, ➀, ➁ and ➂ in Figure 2. In the sequel, we will consider the *k*_*W*_ parameters as inputs to the chromatin modification circuit since they can be modulated by TFs external to the circuit (see SI-equations (S.82),(S.83)).

**Figure 2:**
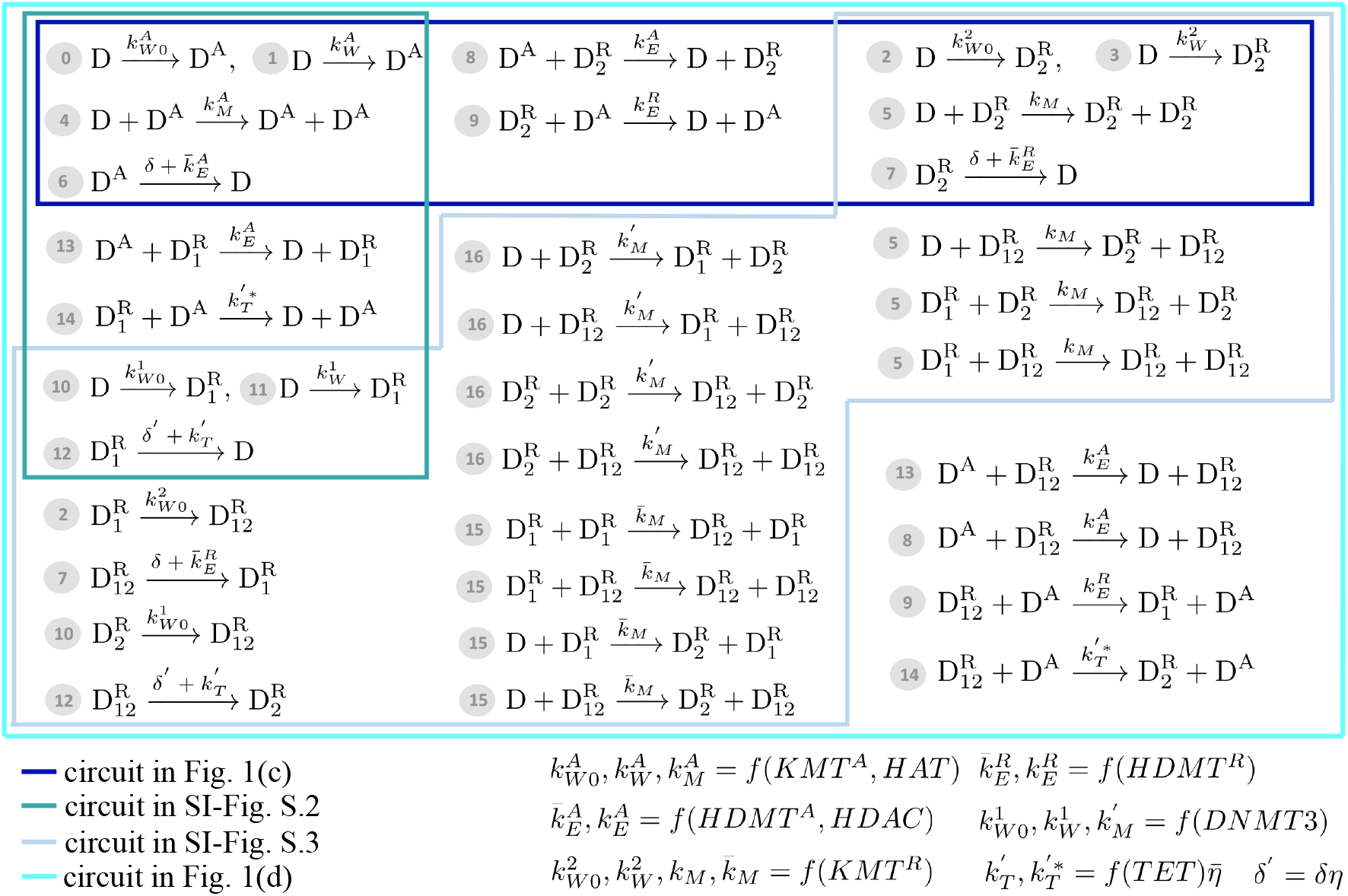
Reactions associated with the chromatin modification circuit motifs. Each reaction is associated with a number, which is referred to in the main text. Specifically, reactions ⓪, ➀, ➁, ➂, ➉ and ⑪ describe *de novo establishment*. Reactions ➃ and ➄ describe *auto-catalysis*, wherein a modification recruits writers of the same modification to nearby nucleosomes. Reactions ⑮ and ⑯ describe *cross-catalysis*, wherein DNA methylation recruits writers of repressive histone modifications and *viceversa*, respectively. Reactions ➅, ➆, and ⑫ represent *basal erasure* while reactions ➇, ➈, ⑬, and ⑭ represent *recruited erasure*, wherein competing modifications recruit erasers of each other. The different colored lines delimit the sets of reactions that take place for each of the circuit motifs shown in Figure 1 and SI-Figures S.2-S.3. Specifically, SI-Figure S.2 depicts the circuit between activating histone modifications and DNA methylation and SI-Figures S.3 depicts the circuit among repressive modifications. Here, we use one-step enzymatic reaction models [68] to capture histone and DNA modifications. In the bottom-right corner, we indicate how the reaction rate constants depend on the amount of writers and erasers, in which *f* (·) is a generic increasing function. Here, *η* and 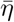 are between 0 and 1 and defined in Section 2.2. Specifically, *η* = 1 in the absence of DNMT1 and 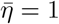 in the absence of MBD (see SI-Section S.1 for the detailed form of the reaction constants and their derivation).

#### Auto-catalysis

Histone modifications can recruit more alike modifications to nearby unmodified histones through a read-write mechanism. Specifically, protein readers bind the modification and recruit writers of the same modification to nearby unmodified histones ([10], Chapter 22 and [36, 37, 27, 38]). For example, protein HP1*α* contains a chromodomain that specifically recognizes H3K9me3 and recruits the Suv39H1 methyltransferase responsible for H3K9 methylation [36]. Similarly, WDR5 is a protein reader that recognizes specifically H3K4me3 and binds to SETs to recruit them for H3K4 methylation [27]. Similarly, for histone acetylation, the p300/CBP complex is a HAT that also has a bromodomain, so it is a reader and a writer at the same time ([10], Chapters 4, 21 and [5], Chapter 2). We capture this auto-catalytic process by reactions ➃ for activating histone modifications and by reactions ➄ for repressive histone modifications in Figure 2.

#### Recruited and basal erasure

An activating (repressing) histone modification can recruit erasers for a repressing (activating) histone modification [39]. For example, JMJD2A is an eraser for H3K9me3/2 and de-methylates H3K9me3/2 through its Jumonji domain while being able to bind H3K4me3 through the Tudor domain. Therefore, H3K4me3 helps recruit this H3K9me3 eraser to DNA. In turn, JARID is an eraser of H3K4me3 and does so through one of its PHD domains. Through a different PHD domain, it binds H3K9me3. Therefore, H3K9me3 helps recruit this H3K4me3 eraser to DNA. Furthermore, the CHD4 subunit of the NuRD (nucleosome remodeling and de-acetylase) complex contains a domain (a PHD domain) that recognizes H3K9me3. This provides a mechanism through which H3K9me3 recruits HDACs. The corresponding reactions are number ➇ and ➈ in Figure 2. Furthermore, in addition to recruited erasure, histone modifications are subject to basal erasure, wherein a modification can be removed passively through dilution due to DNA replication during S phase or through non-specific de-methylation ([10], Chapter 22). Basal erasure is captured by reactions ➅ and ➆ in Figure 2. The resulting circuit diagram is shown in Figure 1(c), wherein each modification auto-catalyzes itself and inhibits the other one.

### 2.2 DNA methylation and crosstalk with histone modifications

Similar to histone modifications, DNA methylation can be captured by enzymatic reactions. Specifically, *de novo* establishment of CpG methylation is mediated by the DNMT3 enzyme (reactions ➉ and ⑪ in Figure 2) and copy-write occurs through the DNMT1 enzyme, which quickly copies the methylation pattern on the nascent DNA strand at replication. De-methylation occurs passively through dilution due to DNA replication and by an active de-methylation mediated by TET enzymes ([10], Chapter 15). In particular, TET enzymes recognize CpGme dinucleotides and have catalytic activity to convert methylated CpG to hydroxilmethylated CpG, then to formylcytosine, and finally to carbolxylcytosine [40, 41]. None of these modified forms are recognized by DNMT1 and therefore they are subject to dilution *via* DNA replication ([5] Chapter 17). The ability of TET enzymes to convert methylated CpG to hydorxilmethylated CpG *in vivo* is hampered by the binding of MBD proteins to methylated CpG [42]. Reversely, MBD proteins cannot bind hydroxylmethylated DNA [41]. The combination of these processes can be captured by a basal erasure reaction (reaction ⑫ in Figure 2), where the expressions of the decay rate constants *δ*′ and 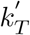 are derived in SI-Section S.1.3, SI-equations (S.33),(S.62). Specifically, we have that *δ*′ = *δη*, in which *δ* is the dilution rate constant, *η* = 0 if the efficiency of the maintenance process by DNMT1 is 100%, and *η* = 1 if DNMT1 is absent, consistent with earlier models [43]. Similarly, we have that 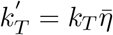, in which *k*_*T*_ is a constant proportional to the level of TET enzyme, 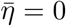 if MBD proteins are highly abundant, and 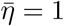 if MBD proteins are absent.

#### Recruited erasure between activating histone modifications and DNA methylation

DNA methylation and H3K4 methylation/acetylation have mechanisms to recruit erasers of each other. Specifically, methylated CpGs recruit MeCP2 proteins, which associate with HDACs to establish de-acetylation ([10], Chapter 15 and [44, 45, 46]). Similarly, methylated CpGs recruit MBD2, which interacts with the NuRD complex to also promote de-acetylation ([44], [10], Chapter 21). These processes are captured by reaction ⑬ in Figure 2. On the other hand, TET1 enzyme has high propensity to bind to unmethylated CpGs through the CXXC domain ([5], Chapter 17 and [41]). This suggests a potential mechanism by which methylated H3K4recruits TET1 to nearby methylated CpGs, enhancing their de-methylation. This is captured by reaction ⑭ in Figure 2, in which the reaction constant 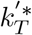 has the same trend as 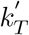 with the abundance of MBD (SI-equation (S.76)). A diagram representing the mutual inhibition between activating histone modifications and DNA methylation is shown in SI-Figure S.2.

#### Cross-catalysis between repressive chromatin modifications

DNA methylation and H3K9 methylation reinforce each other. Specifically, MBD proteins recognize methylated CpG dinucleotides and recruit both histone modifying complexes to the methylated sites. MBD1, in particular, binds to methylated CpG sites and recruits histone methyltransferases for H3K9, SETDB1 and Suv39H1, which bring H3K9me3 about [44]. Similarly, MeCP2 binds methylated CpGs and recruits histone methylases that lead to H3K9me3 [47]. We capture the process by which DNA methylation recruits H3K9 methyltransferases by reactions ⑮ in Figure 2. On the other hand, DNMT3 binds to HP1 protein (reader of H3K9me3), suggesting that H3K9me3 recruits DNA methylation enzymes through HP1 [48]. This is captured by reactions ⑯ in Figure 2. A diagram representing these synergistic interactions is shown in SI-Figure S.3. These interactions create positive feedback loops, wherein each modification enhances the creation of the other.

By combining the competitive interactions between activating and repressive modifications (Figure 1(c) and SI-Figure S.2), with the cross-catalysis among repressive modifications (SI-Figure S.3), we obtain the full chromatin modification circuit of Figure 1(d), whose corresponding reactions are listed in Figure 2. SI-Sections S.1.1 - S.1.7 contain the derivation of the models and of reaction constants. The expressions of reaction constants are function of the level of eraser enzymes, writer enzymes, methyl-DNA binding proteins, and of cell division rate. Figure 2 summarizes these dependencies in a qualitative way. SI-Section S.1.8 contains the list of assumptions that were made to write these reactions. In particular, in this model we do not include spatial regulation for simplicity, assuming that the higher-order structure of chromatin allows any modified nucleosome to act on any other nucleosome within the gene, as in previous work [26]. Even if the model does not include spatial regulation, it still includes “beyond-neighbor” interactions that play a role in bistability [26].

## 3 Results

We analyze both the deterministic and stochastic behavior of the chromatin modification circuit model, focusing on the question of temporal duration of memory of transient input stimuli. Therefore, we regard this circuit as a dynamical system that takes 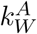 and 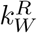, modulated by the binding of sequence-specific TFs to DNA, as inputs and gives as output the chromatin state, captured by the number of modified nucleosomes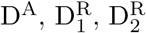, and 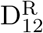 within the gene (Figure 3(a)-(b)).

**Figure 3:**
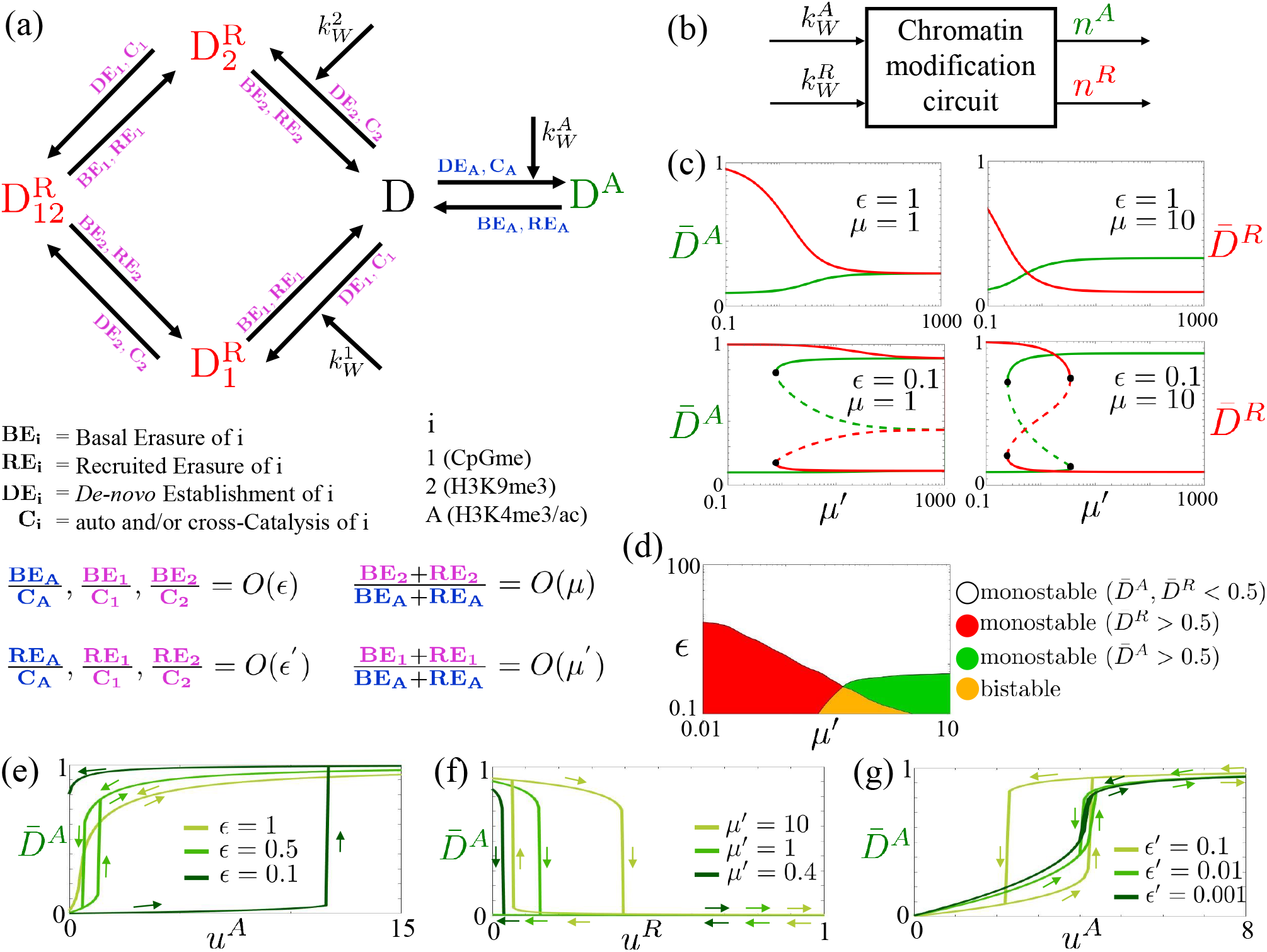
Time scale parameters *ϵ, ϵ*′, *µ* and *µ*′ control bistability and hysteresis in the chromatin modification circuit. (a) Diagram of the gene’s inner chromatin modification circuit in which, compared to Figure (d),we removed the lines and arrows indicating recruitment and catalysis. The labels on each arrow specify the processes enabling that nucleosome modification as indicated below the reaction diagram. We use purple labels for repressive modifications and blue labels for activating modifications. A visual representation of the relationships between the rates of these processes and the parameters *ϵ, ϵ*′, *µ* and *µ*′ is provided. In particular, *ϵ* and *ϵ*′ quantify the time scales of basal and recruited erasure rates of all modifications relative to those of auto and cross-catalysis. Similarly, *µ* and *µ*′ quantify the time scales of erasure rates (basal and recruited) of repressive histone modifications and DNA methylation, respectively, relative to those of activating histone modifications. For the mathematical definition, refer to equation (1) and the related text. (b) Block diagram corresponding to the chromatin modification circuit. Here, 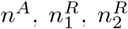 and 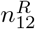 denote the numbers of modified nucleosomes 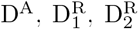, and 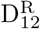 within the gene, 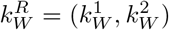 and 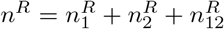. The pair 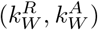 are the inputs and (*n*^*R*^, *n*^*A*^) are the outputs. (c) Steady states of the system as a function of *ϵ, µ* and *µ*′. Here, 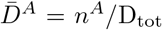 and 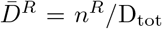 are the fractions of nucleosomes with activating or repressive modifications within the gene with a total of D_tot_ nucleosomes. Plots are obtained from system (2) with 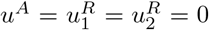. The solid lines represent stable steady states, the dashed lines represent unstable steady states and the black circle represents the bifurcation point (saddle-node bifurcation). In these plots 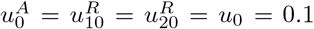 and all the other parameters are set equal to 1 (SI-Figure S.11 shows different values). (d) Chart depicting the (*ϵ, µ*′) combinations that result in a monostable (red, green or white) or bistable (yellow) system for *µ* = 10 (SI-Figure S.12 shows different values of *µ*). Here, *ϵ*′ = 1. (e) Input/output steady state characteristics displaying hysteresis for the 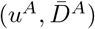 and 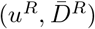 pairs, with 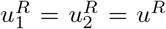, for different values of *ϵ* obtained from simulations of system (2). We consider 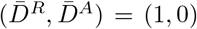 as initial conditions and we set 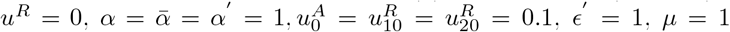, and *µ*′ = 0.8 (SI-Figures S.13 show different values of *µ, µ*′ and *ϵ*′). (f) Input/output steady state characteristics for the 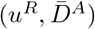 pair, for different values of *µ*′ obtained from simulations of system (2). We consider 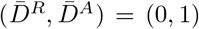 as initial conditions and we set 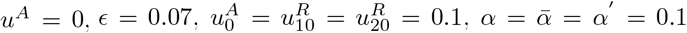, and all the other parameters equal to 1 (SI-Figure S.15(a) shows the 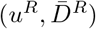 steady state characteristics for the same parameter values). (g) Input/output steady state characteristics for the 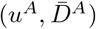 pair, for different values of *ϵ*′ obtained from simulations of system (2). We consider 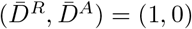 as initial conditions and we set 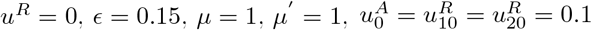 and all the other parameters equal to 1 (SI-Figure S.15(b) shows the 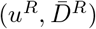 steady state characteristics for different values of *ϵ*′). In all plots 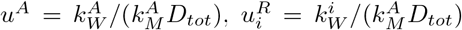 for 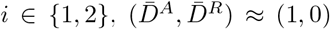 corresponds to the active state and 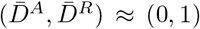 corresponds to the repressed state. In the figure, we use green and red, respectively, to indicate the activating and repressive modifications and related quantities.

We then consider two applications: TF-enabled positive autoregulation for robustness of the active gene state to undesirable repressive input stimuli, such as due to endogenous silencing, and mutually repressing positively autoregulated chromatin modification circuits for long-term persistence of multiple distinct gene expression patterns.

### 3.1 Time scale separation among constituent processes control bistability, hysteresis, and time to memory loss

We first analyze the ordinary differential equation (ODE) model of the chromatin modification circuit, focusing on how parameters affect bistability and hysteresis. Bistability is the co-existence of two stable steady states, specifically active and repressed chromatin states, in the absence of an input stimulus. Hysteresis occurs when the output of the system follows two different paths depending on whether the input is increased or decreased. In particular, the output remains in the proximity of the value reached for high input when this is removed that is, the system “remembers” a transient input stimulus. We then consider a stochastic model to address the question of temporal duration of this memory, given that noise intrinsic to molecular reactions can always displace the system’s state.

For our analysis, we let 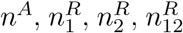 and *n*^*D*^ denote the number of nucleosomes with activating (D^A^), repressive (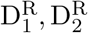, and 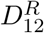), and no modifications (D), respectively, within a gene composed of a total of D_tot_ nucleosomes. All analyses are performed considering the unitless normalized time 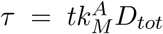 with *D*_*tot*_ = D_tot_*/*Ω and Ω the reaction volume. We define the following non-dimensional parameters, capturing the characteristic time scales of the circuit’s constituent processes:

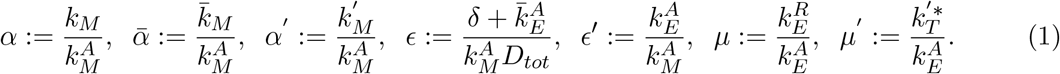

Here, *α* is the non-dimensional auto-catalysis rate parameter, while 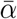, and *α*′ are the non-dimensional cross-catalysis rate parameters, which we assume on the same order without loss of generality. We further let 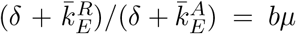 with *b* = *O*(1), implying that *µ* scales the ratio between the erasure rates of repressive and activating histone modifications. We similarly let 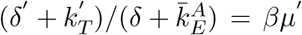 with *β* = *O*(1), implying that *µ*′ scales the ratio between the erasure rates of DNA methylation and those of activating histone modifications. From these relationships, it also follows that 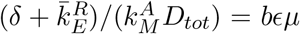 and that 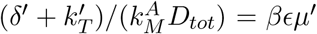. Therefore, the parameter *ϵ* scales the ratio between the basal rate at which all modifications are erased and the rate at which they are auto or cross-catalyzed. Finally, since 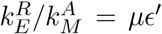 and 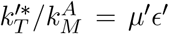, it follows that *ϵ*′ scales the ratio between the rate at which modifications are removed through mutual recruitment of erasers and that at which they are auto or cross-catalyzed. These parameter definitions are pictorially conveyed in Figure 3(a).

Putting these definitions together, in our model, *ϵ* and *ϵ*′ are non-dimensional parameters that quantify the time scales of basal and recruited erasure rates of all modifications relative to those of auto and cross-catalysis. Similarly, *µ* and *µ*′ are non-dimensional parameters that quantify the time scales of erasure rates (basal and recruited) of repressive histone modifications and DNA methylation, respectively, relative to those of activating histone modifications. Since these normalized parameters are functions of the biochemical reaction rate constants (equation (1)) and, in turn, the reaction rate constants are known functions of the level of writer and eraser enzymes, TET enzymes, and MBD proteins (see Figure 2 and SI-Section S.1.6), it is possible to experimentally vary these parameters by over-expressing, inhibiting, or by recruiting these molecular players.

#### Deterministic behavior

We first analyze the ordinary differential equation (ODE) model of the chromatin modification circuit, focusing on how the above parameters affect dynamics. To this end, we assume that the number of total modifiable units D_tot_ is sufficiently large such that the variables 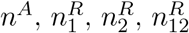 and *n*^*D*^ can be considered real-valued and thus their temporal evolution can be described by ODEs. In particular, we describe the system in terms of fractions of modifiable units, that is 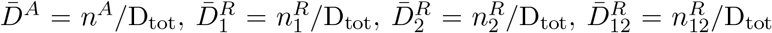 and 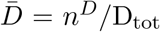. We further introduce the normalized inputs 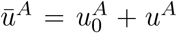 with 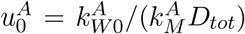 and 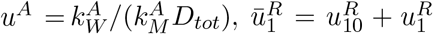 with 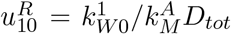 and 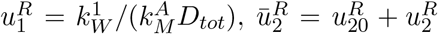 with 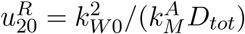 and 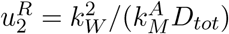. With these definitions and letting 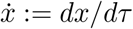, the ODEs describing the system in non-dimensional variables are given by

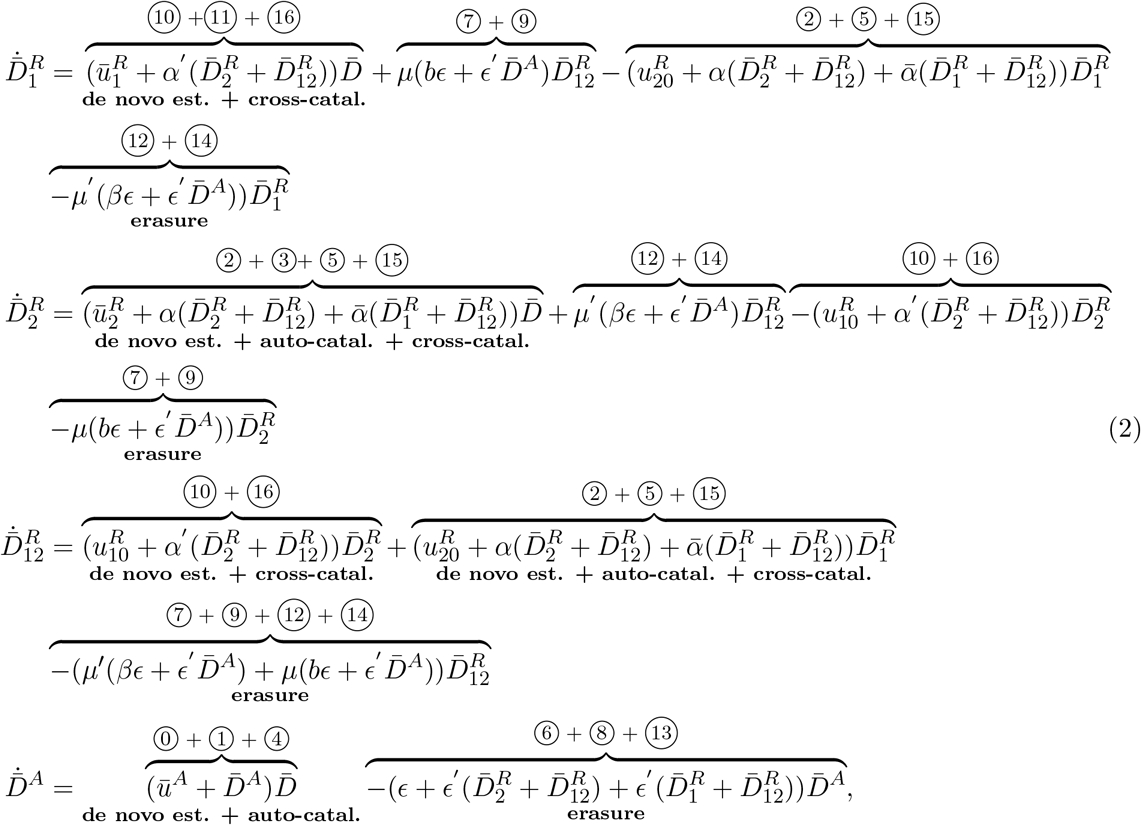

in which 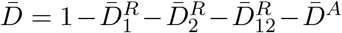 and the circled numbers indicate the corresponding reactions from Figure 2 (see SI-Section S.1.6 and SI-equations (S.80) for derivation).

In the absence of inputs 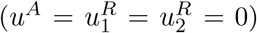, system (2) has a unique stable steady state, if *ϵ* is not sufficiently small. This steady state has 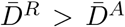 for small *µ*′. When *ϵ* is sufficiently small, the system is bistable, with both active 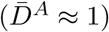 and repressed 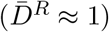 co-existing stable chromatin states. But when *µ*′ is very small, even for small *ϵ*, the system returns monostable with a unique stable steady state corresponding to the repressed state (Figure 3(c)-(d) and SI-Figure S.11). Furthermore, as *µ*′ increases (DNA methylation erasure rate increases), the system approximates well the histone modification system (Figure 1(c)), wherein DNA methylation is not present. This is because when *µ*′ is sufficiently large, the DNA methylation is erased very quickly and then 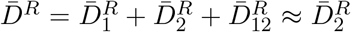. As a consequence, when *ϵ* is sufficiently small the system is bistable for intermediate values of *µ*, monostable with 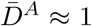 when *µ* is large and monostable with 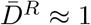 when *µ* is small (SI-Figure S.4). Varying *ϵ*′ does not change these trends, except for *ϵ*′ ≪ *ϵ*, in which case the system returns monostable (SI-Figure S.11).

Figure 3(e) shows the input/output steady state characteristics of the chromatin modification circuit. Specifically, as *u*^*A*^ is increased starting from 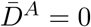, the state switches to 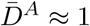 for *u*^*A*^ above a critical value, which increases as *ϵ* decreases. When *u*^*A*^ is decreased back to zero, the path that 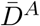 follows is different and it can keep 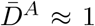 even when *u*^*A*^ = 0 if *ϵ* is sufficiently small. In this case, we say that the state transition becomes irreversible and *the system keeps memory* of its input stimulus. From a biological point of view, this implies that an activating transient input stimulus allows to permanently re-activate a silenced gene. This memory vanishes as *ϵ* increases. When *µ*′ decreases, the value of *u*^*A*^ required to flip the system’s state to fully active increases until we reach a regime under which the memory is lost (SI-Figure S.13). A similar behavior is observed for the relationship between the repressing input *u*^*R*^ (with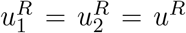) and 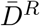 (SI-Figure S.14). Biologically, this implies that a repressive transient input stimulus can permanently silence an initially active gene. However, as opposed to the 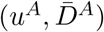 characteristics, if we reduce *µ*′, the minimum value of *u*^*R*^ required to silence the system 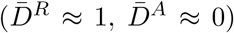 decreases (Figure 3(f), SI-Figure S.14). This also shows that as *µ*′ is decreased, the active state becomes less robust to repressive input stimuli.

A comparison of the plots of SI-Figure S.13 and SI-Figure S.14 for small *ϵ* and *µ* = 1 reveals that, even when *µ*′ = 1, it takes smaller values of *u*^*R*^ to switch the state to 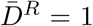 than the values of *u*^*A*^ required to switch the state to 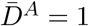. This fact highlights that it is the structure itself of the chromatin modification circuit, rather than the rate constants alone, to “bias” the system towards a repressed chromatin state. In fact, the two-layer topology of the repressive modifications confers increased robustness to the repressed state with respect to activating stimuli 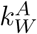 (Figure 3(a)). This is in contrast to what is observed for the system that does not include DNA methylation (Figure 1(c)), wherein once all parameters are equal to each other, the stability landscape is symmetric. (SI-Section S.2.1). Varying *ϵ*′ does not significantly affect these trends (SI-Figures S.13-S.14), unless *ϵ*′ ≪ *ϵ*, in which case hysteresis is lost (Figure 3(g)).

In summary, when recruited erasure (*ϵ*′) and auto-catalysis/cross-catalysis 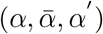 are sufficiently faster than basal erasure (*ϵ*), the circuit displays bistability and hysteresis, except if *µ* and/or *µ*′ are sufficiently small. In this case, the unique stable state is the fully repressed chromatin state, independent of all other parameters.

#### Stochastic behavior

We next evaluate how the stationary distribution of the chromatin state, the time to memory loss, and the ability to reactivate a repressed gene depend on *ϵ, ϵ*′, *µ* and *µ*′, and on the inputs *u*^*A*^ and *u*^*R*^. To this end, we first perform a computational study by using Gillespie’s Stochastic Simulation Algorithm (SSA) [49]. We then complement this with a mathematical derivation of the approximate expressions of both the stationary distribution and the time to memory loss as a function of the parameters and input stimuli.

Figure 4(a)-(b) shows how the stationary distribution is shaped by *ϵ* and *µ*′ (a) and by inputs *u*^*R*^ and *u*^*A*^ (b). Specifically, *ϵ* has a most critical role. As it decreases, with all other parameters remaining fixed, the peaks of the distribution become more concentrated about the fully repressed and fully active chromatin states. Highly concentrated peaks imply that the probability of finding the chromatin state outside of these configurations is very small. This property is connected to the *time to memory loss*, defined as the expected value of the earliest time the chromatin state reaches the active state starting from the repressed state and *viceversa*. In fact, as *ϵ* is decreased, the chromatin state starting in either the repressed or active state takes longer, on average, to reach the active or repressed state for the first time, respectively (Figure 4(c)-(d)). In contrast to what found for the histone modification model of Figure 1(c) (SI-Figure S.6(a)), decreasing *ϵ* also has an effect in increasing the height of the peak in correspondence of the repressed state to the detriment of the peak corresponding to the active state (Figure 4(a)). When *µ*′ is decreased, this bias increases, and the peak in correspondence of the repressed state becomes more concentrated (Figure 4(a)). Varying *µ* has a similar effect as varying *µ*′ (SI-Figure S.19). Inputs *u*^*R*^ and *u*^*A*^ each increase the height of the peak corresponding to the repressed and active state, respectively (Figure 4(b)). Consistent with the deterministic analysis (SI-Figure S.13), for smaller *µ*′, it is required a larger *u*^*A*^ to increase the probability of finding the state in the fully active chromatin state (SI-Figure S.18(b)). By contrast, *ϵ*′ does not significantly affect the trends observed for *ϵ, µ*′, *µ* and the inputs, although it modulates some aspects of the distribution (SI-Section S.3.5 and SI-Figures S.17-S.20). Specifically, when *ϵ*′ decreases compared to *ϵ*, the peaks of the distribution become less concentrated and, in the extreme case where *ϵ*′ ≪ *ϵ*, the distribution becomes unimodal (Figure 4(e)).

**Figure 4:**
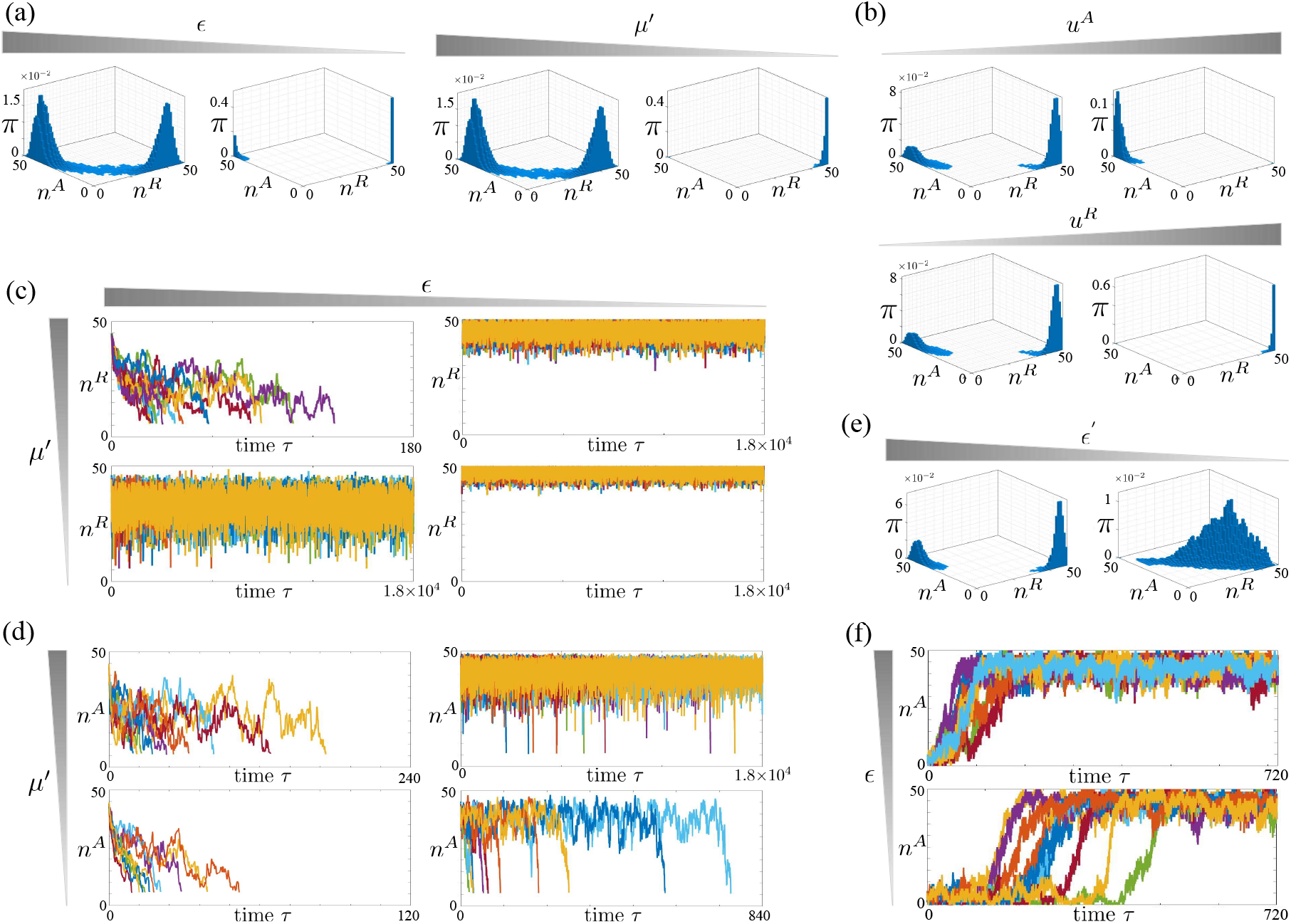
Time scale separation parameters *ϵ, ϵ*′ and *µ*′ control bimodality, time to memory loss, and reactivation time in the chromatin modification circuit. (a)-(b) Stationary probability distributions *π* of (*n*^*A*^, *n*^*R*^) for the chromatin modification circuit in Figure 3(a) obtained by simulating the reactions listed in Figure 2 with the SSA. (a) Effect of *ϵ* and *µ*′ on the distribution: in the left side plots *ϵ* = 0.19, 0.02 and *µ*′ = 1. In the right side plots, *µ*′ = 1, 0.1 and *ϵ* = 0.19. (b) Effect of the input stimulus on the distribution: here, *ϵ* = 0.12 and *µ*′ = 0.8. In the top plots, *u*^*A*^ = 0, 1, and in the bottom plots 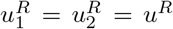 and *u*^*R*^ = 0, 1 (SI-Figure S.18 show different values of *µ*′ and *ϵ*). The parameter values for each panel are listed in SI-Table 1. In all simulations, *ϵ*′ = 1 and *µ* = 1 (SI-Figures S.19-S.20 show different values) and we decrease *ϵ* by decreasing 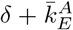 (similar results can be obtained if we vary 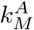 as shown in SI-Figure S.17). (c) Time trajectories of *n*^*R*^ starting from a repressed chromatin state 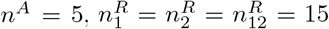, as *ϵ* and *µ*′ are varied. Simulations are stopped when *n*^*R*^ = 6 for the first time. (d) Time trajectories of *n*^*A*^ starting from an active chromatin state 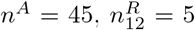, as *ϵ* and *µ*′ are varied. Simulations are stopped when *n*^*A*^ = 6 for the first time. In all plots of (c)-(d), time is normalized according to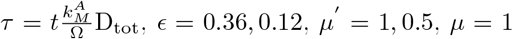, and *ϵ*′ = 1 (SI-Figure S.22 shows different values of *ϵ*′). The parameter values are listed in SI-Table 2. (e) Effect of *ϵ*′ on the stationary probability distribution *π*. We set *ϵ* = 0.12, *µ*′ = 1, *µ* = 1 and *ϵ*′ = 1, 0.001 from left to right (SI-Figure S.21 shows different values of *ϵ*′). The parameter values for each panel are listed in SI-Table 24. (f) Time trajectories of the system starting from *n*^*R*^ = 45, *n*^*A*^ = 5 and applying an input *u*^*A*^ that, at steady state, leads to a unimodal distribution in the proximity of the active state *n*^*A*^ = D_tot_. The parameter values are listed in SI-Table 3. In particular, *u*^*A*^ = 3.2, *µ*′ = 0.1, *ϵ* = 0.24, 0.16, *µ* = 1 and *ϵ*′ = 1 (SI-Figure S.23 shows different parameter values). In all plots, 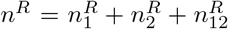 and simulations are obtained by implementing the set of reactions listed in Figure 2 with the SSA [49]. In all simulations, 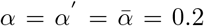. In our model, parameters *ϵ* and *ϵ*′ quantify the time scales of basal and recruited erasure rates of all modifications relative to those of auto and cross-catalysis. Similarly, parameters *µ* and *µ*′ quantify the time scales of erasure rates (basal and recruited) of repressive histone modifications and DNA methylation, respectively, relative to those of activating histone modifications. Mathematical definitions are found in equation (1). In all plots (*n*^*A*^, *n*^*R*^) ≈ (50, 0) corresponds to the active state and (*n*^*A*^, *n*^*R*^) ≈ (0, 50) corresponds to the repressed state. In each panel of (c),(d), (f), the number of trajectories plotted is 10.

In the regime where both *ϵ*′ ≪ 1 and *ϵ* ≪ 1, in which the system displays a bimodal distribution (SI-Figures S.17-S.20), we can analytically derive a one-dimensional Markov chain approximation that allows analytical computation of the stationary distribution and of the time to memory loss as functions of the parameters (SI-Sections S.3.2-S.3.4). Biophysically, in this regime, the reactions represented by the label BE_i_ in Figure 3(a) become slower compared to those represented by the label C_i_. In particular, letting 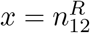 with *x* ∈ [0, D_tot_], we have that, when *ϵ* ≪ 1, the stationary probability distribution *π*(*x*) can be approximated by

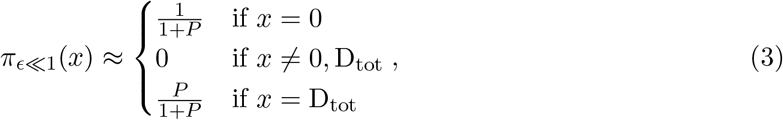

with

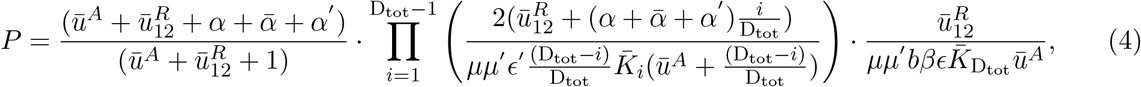

in which 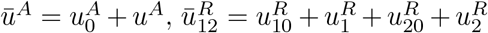 and 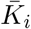, defined in SI-equations (S.161),(S.171), independent of *ϵ, µ, µ*′ and *ϵ*′ (see SI-Section S.3.3 for the mathematical derivations). From this expression, we conclude that if *ϵ* ≪ 1, *π*(*x*) ≈ 0 for all *x* except for *x* = D_tot_ (fully repressed state) and *x* = 0 (fully active state), implying that the probability of finding the system in one of the intermediate states is about zero. This is in accordance with the computational results of Figure 4(a), which show higher concentration around the extreme states, and with Figure 4(c)-(d), which show smaller likelihood of transitioning out of those extreme states as *ϵ* is decreased. When *ϵ* goes to zero, *P* → ∞, which implies that *π*(D_tot_) ≈ 1. This is consistent with the structural bias of the chromatin modification circuit towards the repressed chromatin state (Figure 3(a)) and the computational results of SI-Figure S.18(a). This bias is enhanced as *µµ*′ is decreased since *P* → ∞ as *µµ*′→ 0, leading to *π*(D_tot_) ≈1. Furthermore, in agreement with what observed in Figure 4(b), an increase of *u*^*A*^ decreases *P* and hence increases *π*(0). By contrast, an increase of *u*^*R*^ increases *P*, which, in turn, increases *π*(D_tot_). Also, larger *u*^*A*^ values are required to increase the probability of the active state to the same level when *µ*′ is smaller.

Similarly, the normalized time to memory loss of the repressed state can be approximated as (SI-Section S.3.4)

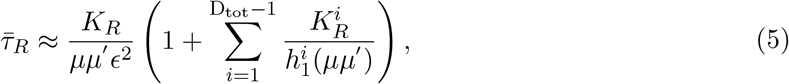

with 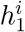 an increasing function, 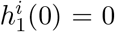, and *K*_*R*_ and 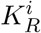 functions independent of *ϵ, µ*′ and *µ*, while the normalized time to memory loss of the active state can be approximated as

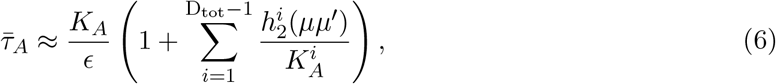

with 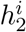 an increasing function and 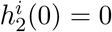, and *K*_*A*_ and 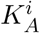 functions independent of *ϵ, µ*′ and *µ*. In the limiting condition where *ϵ* → 0, we have that both 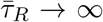 and 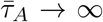. Therefore, decreased *ϵ* is the driver for increasing the extent of epigenetic memory of both the active and repressed chromatin states. Furthermore, because of the structural asymmetry of the chromatin modification circuit, the power of *ϵ* is different in the two above expressions, indicating that decreased *ϵ* increases the extent of memory much more for the repressed state than for the active state. This is in contrast with what obtained for the histone modification circuit, in which both 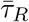 and 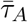 are *O*(1*/ϵ*) (SI-equations (S.138) and (S.140)). Finally, decreased *µ* and *µ*′ lead to larger 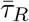 but to lower 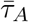. These trends are in agreement with the simulations in Figure 4(c)-(d). In particular, in each panel, we plotted a number of trajectories equal to 10 to provide a comprehensive representation of the parameter effects on the trajectories and, at the same time, a clear reading of the figure.

We next determine *via* simulation how the parameters *ϵ, µ*′ and *ϵ*′ affect the time to reactivation, i.e., the time it takes to re-activate an initially repressed chromatin state by applying a sufficiently large activating input stimulus *u*^*A*^ (Figure 4(f)). The time trajectories show a switch-like behavior, which is more prominent for *ϵ* ≪ 1, *µ*′ ≪ 1, and *ϵ*′ *>* 1 (SI-Figure S.23). Also, the latency, i.e., the time the trajectory takes to switch after an activating input is applied, is highly variable and its variability is mostly dictated by small *ϵ* and small *µ*′, which, together, make gene reactivation a low probability event. Furthermore, the time to reactivation decreases by increasing *µ*′ and/or increasing *ϵ*, consistent with results from the deterministic model (SI-Figure S.24).

Taken together, our results indicate that basal erasure rate sufficiently smaller than recruited erasure rate (*ϵ* sufficiently small and *ϵ*′ sufficiently larger than *ϵ*) allows a bimodal distribution such that both active and repressed chromatin states each have high probabilities compared to intermediate states. As *ϵ* decreases, this distribution is more concentrated about the fully active and repressed states and the time to memory loss of either state increases. Even when there is no asymmetry between activating and repressive modification erasure rates (*µ* = 1 and *µ*′ = 1), the time to memory loss is larger for the repressed chromatin state. This asymmetry is accentuated by *µ*′ *<* 1.

#### Relationship with published data

In [50], the authors study the kinetics of silencing and reactivation of chromatin modifications, including H3K9me3 and DNA methylation, at the single-cell level. In particular, they show that silencing and reactivation has a all-or-none behavior. Our model predicts that this all-or-none response is observed when the underlying chromatin modification circuit has a bimodal stationary distribution, which we proved requires *ϵ* ≪ 1 and *ϵ*′ sufficiently larger than *ϵ* (SI-Figures S.23). This suggests that in practice we should expect *ϵ* ≪ 1 and *ϵ*′ not too small. Furthermore, data on temporal dynamics of DNA de-methylation *in vivo* indicate that *µ*′ ≪ 1 (SI-Section S.3.1). Early elegant experimental studies performed by [51] further show that the latency of reactivation of a silenced gene is highly variable. In our model, a highly variable latency of reactivation events is predicted under the parameter regime where both *ϵ* and *µ*′ are small (Figure 4(f) and SI-Figure S.23), suggesting that *µ*′ ≪ 1 may in fact be responsible for the experimentally observed latency of reactivation [51]. Furthermore, in [51] it is shown that increased proliferation rate leads to a faster reactivation kinetics. In our model, increased proliferation is captured by increased *ϵ* and in fact higher *ϵ* leads to faster kinetics of reactivation events (SI-Figure S.25(a)-(c)), in agreement with the observations in [51].

Finally, a basic DNA methylation model that does not include MBD proteins cannot reflect the low effective active erasure rates of DNA methylation through the action of TET enzymes, *k*_*T*_, encountered *in vivo* (SI-reactions S.45). In our model, the ability of TET enzymes to convert methylated DNA to hydorxilmethylated DNA is hampered by the binding of MBD proteins to methylated DNA (SI-Section S.1.3). As a result, the effective active erasure rate constant, 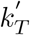, derived in SI-equation (S.62), can be written as 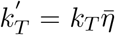 in which *k*_*T*_ is a constant proportional to the level of TET enzyme, where 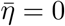 if MBD proteins are highly abundant, and 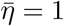 if MBD proteins are absent (SI-equation S.62). This implies that the parameter 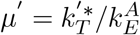 is proportional to the amount of TET enzymes and decreases with the amount of MBD proteins. Lower levels of MBD proteins result in higher *µ*′ and in our model this leads to faster reactivation kinetics and decreased variability of reactivation time (SI-Figure S.25(d)-(f)). This is consistent with the experimental observations of [52], which show that knocking down of MBD proteins not only makes the reprogramming process faster, with almost the entire cell population converted into iPS cells within 7 days, but makes it also almost deterministic.

### 3.2 Model of transcriptional regulation by chromatin state

Chromatin state affects transcription by modulating nucleosome compaction. DNA wrapped around highly compacted nucleosomes is difficult to access by TFs and therefore genes in such DNA regions are not transcribed. By contrast, DNA wrapped around less compacted nucleosomes is more likely to be accessible by TFs, and hence is susceptible to transcription. Repressive histone modifications and DNA methylation generally lead to increased compaction while activating histone modifications keep a more open chromatin structure ([10], Chapter 3).

We therefore assume that transcription is allowed only by D^A^, but not by D or any of 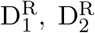, and 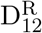. In principle, nucleosome state D may allow some low basal transcription, through non-specific targeting by chromatin remodelers ([53],[32]), but we assume this is sufficiently low to be negligible. A more general model could include this basal rate, accounting for the fact that transcription by RNA Pol II of D occurs concurrently with the deposition of H3K4me3 and hence with the conversion of D to D^A^ [33]. Thus, by lumping together transcription and translation, we obtain the following gene expression model (SI-Section S.4):

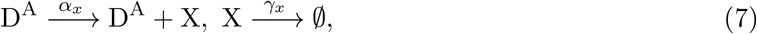

in which X is the gene product, *γ*_*x*_ is the decay rate constant of X and *α*_*x*_ = *F* (*A, R*) is an increasing function of the concentration of TF activators A and a decreasing function of the concentration of TF repressors R. Here, for simplicity, we assume that *α*_*x*_ is a constant. The main conclusions of this paper are not affected by this assumption. For the deterministic analysis, considering again the normalized time 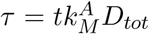, and letting 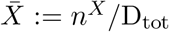, with *n*^*X*^ the number of molecules of X, 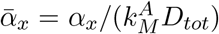, and 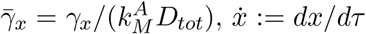, the ODE associated with gene expression in non-dimensional variables can be written as

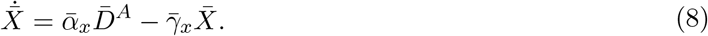

### 3.3 Positive TF-enabled autoregulation extends memory of active chromatin state robustly to perturbations

The study of the chromatin modification circuit reported in the previous section shows that the two-layer topology of the repressive modifications biases the system towards a repressed chromatin state, and that this bias becomes even more pronounced for low values of *µ*′. It follows that, when the chromatin state is in the active configuration, without any activating input stimulus, even a small perturbation can cause a quick silencing of the gene. For an engineered gene expression system, this silencing will disrupt the intended engineered function of the cell line, as observed in experiments with different cell types [16, 17, 18, 19]. Here, we study how TF-enabled positive autoregulation can alleviate this problem by restoring memory of the active gene state and by making it more robust to endogenous silencing. A possible form of positive autoregulation is when the gene product X acts as a TF activator that recruits writers of activating histone modifications to the gene. This is the case of many TFs involved in cell fate determination, such as Oct4 and Nanog [54]. The analysis of this design thus not only provides a potential engineering solution to the silencing problem of chromosomally integrated gene expression systems, but it also allows for a different understanding of the role of positive autoregulation in natural networks.

In this case, the input 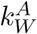 of the chromatin modification circuit is a monotonically increasing function of *n*^*X*^ (SI-Section S.1.7). The positive autoregulation model is thus given by combining the chromatin modification circuit reactions (Figure 2) with the gene expression reactions (7), and the expression for 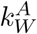 (SI-equation (S.82)). The corresponding interaction diagram is shown in Figure 5(a), and Figure 5(b) shows the block diagram representation with input 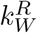 and output *n*^*X*^.

**Figure 5:**
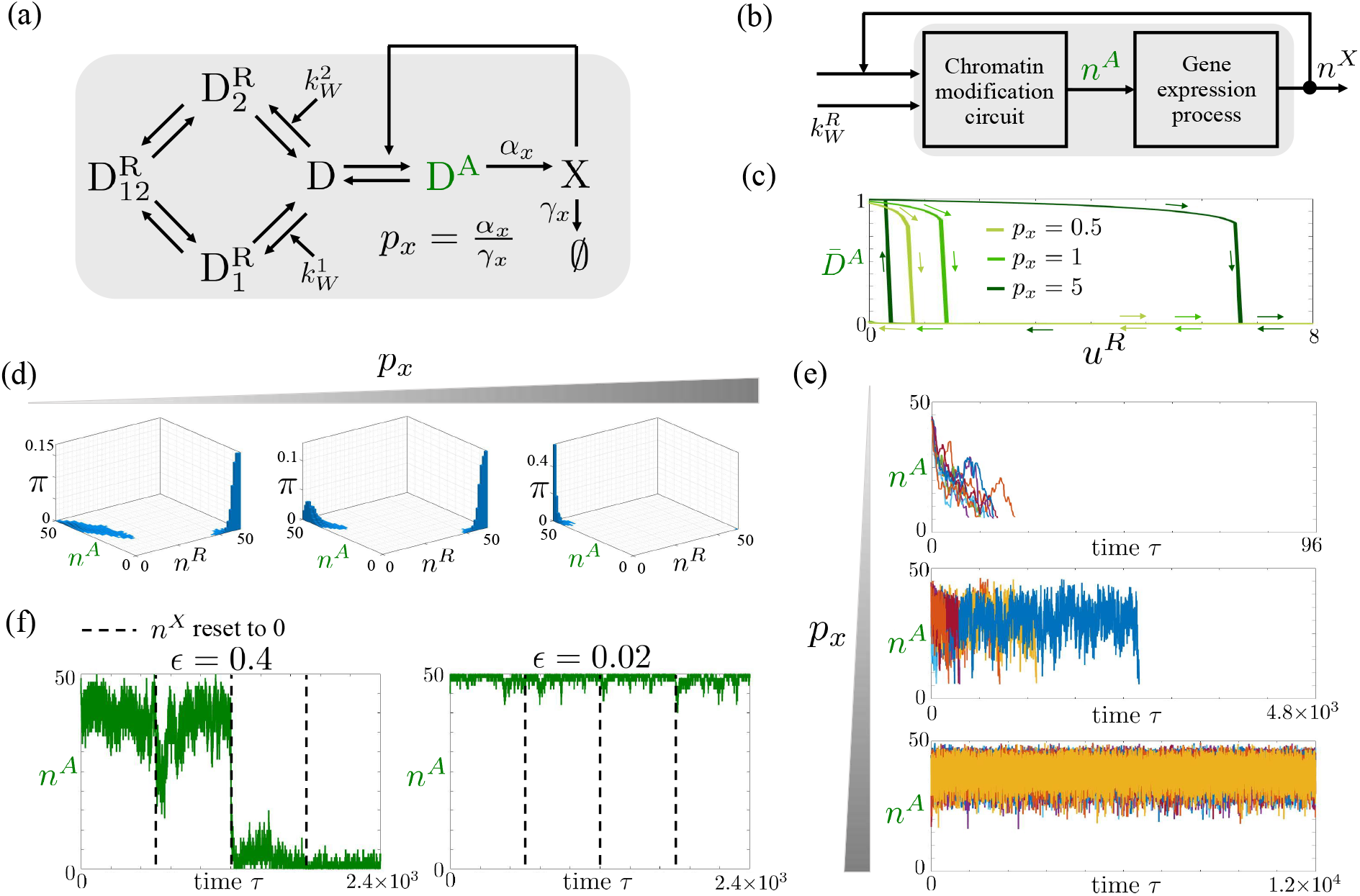
Positive autoregulation allows robust temporal extension of memory of the active chromatin state. (a) Diagram of a positively autoregulated gene where the gene’s product X recruits writers of D^A^. A simplified representation of the chromatin modification circuit is introduced, in which the the labels are not represented. (b) Block diagram corresponding to the circuit in panel (a). Here, 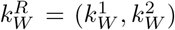. Furthermore, 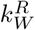 is the input and *n*^*X*^, the number of molecules of X, is the output, which feeds back on 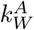 by increasing its value. (c) Input/output steady state characteristics for the 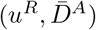 pair, with 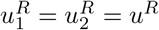, for different values of *p*_*x*_ obtained from simulations of system (S.183) with 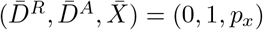 as initial conditions. The parameter values are listed in SI-Table 4. In particular, *ϵ*′ = 1, *µ* = 1, *µ*′ = 0.7, *ϵ* = 0.1. (d) Stationary probability distribution *π* obtained by simulating the reactions listed in SI-Table 5 with the SSA. As before, 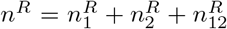. The parameter values of each plot are listed in SI-Table 5. In particular, *p*_*x*_ = 0, 0.1, 10, *ϵ*′ = 1, *µ* = 1, *µ*′ = 0.5, *ϵ* = 0.12 (SI-Figures S.27-S.28 show different parameter values). (e) Time trajectories obtained by simulating the reactions listed in SI-Table 6 with the SSA with no inputs and starting with initial conditions 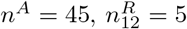 and *n*^*X*^ = *p*_*x*_*n*^*A*^. Simulations are stopped the first time at which *n*^*A*^ = 6. In all plots, time is normalized according to 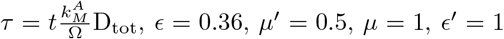, and *p*_*x*_ = 0, 0.2, 5. The parameter values of each panel are listed in SI-Table 6. In each panel, the number of trajectories plotted is 10. (f) Time trajectories of system of reactions listed in SI-Table 5 with the SSA starting from the active chromatin state (*n*^*A*^ = 45, *n*^*D*^ = 5 and *n*^*X*^ = 45). At the indicated times, *n*^*X*^ is reset artificially to zero (dashed lines). The parameter values for each simulation are listed in SI-Table 7. In particular, *p*_*x*_ = 1, *µ*′ = 0.1, *µ* = 1, *ϵ*′ = 1 and *ϵ* = 0.4, 0.02. For both values of *ϵ*, the system is bistable. In our model, *ϵ*, defined in equation (1), is a non-dimensional parameter that quantifies the time scales of basal erasure rate of all modifications relative to those of auto and cross-catalysis. In the figure, we use green to indicate the activating modification and related quantities.

#### Deterministic behavior

By assuming molecular counts sufficiently high to use ODEs and by combining the ODEs of the chromatin modification circuit (2) with those of gene expression (8), we obtain the ODE model of the positive autoregulation system (SI-equations (S.183)). We first realize bifurcation plots with *p*_*x*_ = *α*_*x*_*/γ*_*x*_ the bifurcation parameter (SI-Figure S.26). As *p*_*x*_ increases, the system transitions from monostable with one stable equilibrium point corresponding to the fully repressed chromatin state to bistable in which also a fully active chromatin state is a stable equilibrium. For lower values of *µ*′, higher values of *p*_*x*_ are required to make the fully active state appear as a stable equilibrium. Therefore, increased *p*_*x*_ can compensate, to some extent, for the asymmetry of the system, thus restoring stability of the active state. Varying *µ* has a similar effect as varying *µ*′. In order to gain a qualitative understanding of the way positive autoregulation affects the stability of the system, we also conducted a mathematical analysis assuming that protein dynamics are much faster than chromatin modification dynamics. This analysis shows that positive autoregulation has an effect equivalent to increasing the auto-catalysis rate of the activating histone modification. Based on the bifurcation plots realized for the chromatin modification circuit (SI-Figure S.11), a higher auto-catalysis rate constant 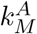 can restore the active state stability, consistent with the results in SI-Figure S.26.

The strength of positive autoregulation *p*_*x*_ also affects the robustness of the active gene state’s memory to endogenous silencing. Specifically, we introduce in our model possible perturbations that can lead to silencing, due to for example chromatin spreading from surrounding gene loci, by considering 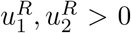. The input/output steady state characteristics 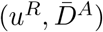 with 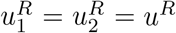, shows persistence of the active state to increasing values of *u*^*R*^ as *p*_*x*_ is increased (Figure 5(c)). These results imply enhanced robustness of the active state to repressive input perturbations.

#### Stochastic behavior

Here, we determine how positive autoregulation modulates the stationary probability distribution of the chromatin state and the time to memory loss of the active state. To this end, we first perform a computational study by simulating the full set of reactions (SI-Table 5) with the SSA [49]. This study shows that, increasing *p*_*x*_, the peak in correspondence to the active state becomes more concentrated and its height increases to the detriment of the peak corresponding to the repressed state (Figure 5(d)). However, for lower values of *µ*′, the height of the peak in correspondence to the active state is also lower, implying that a higher value of *p*_*x*_ is required to increase the peak to the same level (SI-Figure S.28). Reducing *µ* has a similar effect as reducing *µ*′ (SI-Figure S.28). These results are in agreement with the temporal trajectories of *n*^*A*^ (Figure 5(e)), which indicate that the time to memory loss of the active state increases when *p*_*x*_ is increased. The trends with which *p*_*x*_ affects the time to memory loss of the active state do not depend on *ϵ*′ (SI-Figure S.29).

To gain an analytical understanding of how *p*_*x*_ affects the stationary distribution and the time to memory loss, we also computed analytically these quantities as a function of the system parameters in the regime *ϵ*′ ≪ 1 and *ϵ* ≪ 1 and by assuming that protein dynamics are fast compared to chromatin dynamics. The latter assumption may be satisfied if the protein is quickly degrading due to proteolysis [55]. In this regime, we obtain a one-dimensional Markov chain reduction of the system (SI-Section S.5.3), which leads to the stationary probability distribution for *ϵ* ≪ 1 as

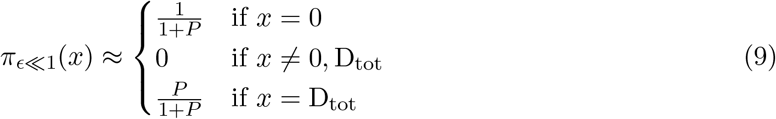

with

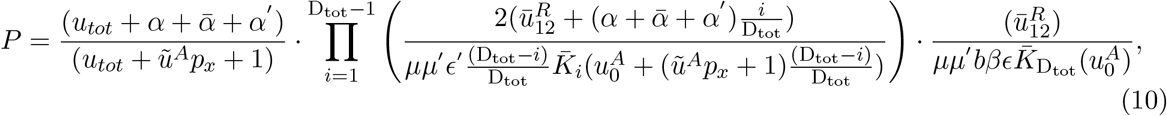

in which 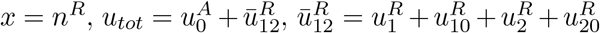 and 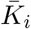 defined in SI-equation (S.161). From these expressions, we conclude that by increasing *p*_*x*_, *P* decreases and, in turn, the height of the peak in correspondence to *x* = 0 (active chromatin state) increases to the detriment of the height of the peak at *x* = D_tot_ (repressed chromatin state). Furthermore, the smaller *µ* and *µ*′ are, the higher *p*_*x*_ has to be in order to raise the peak corresponding to the active state. These results are in agreement with the computational results (Figures 5(d), S.28).

Using the same model reduction, we derive the formula of the normalized time to memory loss of the active chromatin state, which, for *ϵ* ≪ 1, can be approximated as

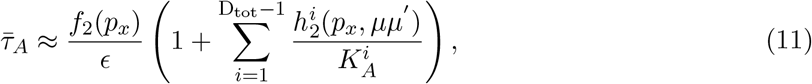

in which *f*_2_ and 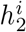 are increasing functions of their arguments, 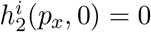, and 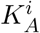 are functions independent of *ϵ, µ*′, *µ* and *p*_*x*_ (SI-Section S.5.4). From this expression, it follows that if *p*_*x*_ increases, 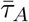 increases. This confirms that positive autoregulation helps extend the memory of the active state.

Taken together, these findings indicate that positive TF-enabled autoregulation contributes to extend the duration of memory of an active chromatin state, thus re-balancing the asymmetry that naturally biases the chromatin modification circuit towards a repressed state. This is also manifested in the effect of positive autoregulation on the time to reactivation of a gene, wherein the time to reactivation decreases by increasing *p*_*x*_ (SI-Figure S.30).

We also determined how the chromatin modification circuit contributes to the robustness of the active state’s memory to perturbations that reduce the availability of X at the gene (Figure 5(f)). For low *ϵ* the robustness is improved, implying that the system is able to keep the active chromatin state even when X is transiently removed. Then, small *ϵ* may be a way in which the chromatin modification circuit enables robustness of the active state’s memory to disruption of the binding of the TF X to DNA, such as due to DNA replication and cell division.

### 3.4 Wiring positively autoregulated chromatin modification circuits allows long-term persistence of gene expression patterns

Long-term and reconfigurable memory of multiple gene expression patterns is desirable for a number of applications, such as programmable cell therapies and *in vivo* transdifferentiation, wherein TFs need to be activated in a specific temporal sequence and in an exclusive fashion [56, 57]. We thus analyze a candidate circuit to implement this function: two positively autoregulated chromatin modification circuits that mutually recruit writers of repressive modifications to each other (Figure 6(a)). This is also a common motif in GRNs involved in cell fate determination, such as the PU.1/GATA.1 antagonism [35, 58], the Nanog/GATA interaction [6], and the Cdx2/Oct4 circuit motif [6, 59].

**Figure 6:**
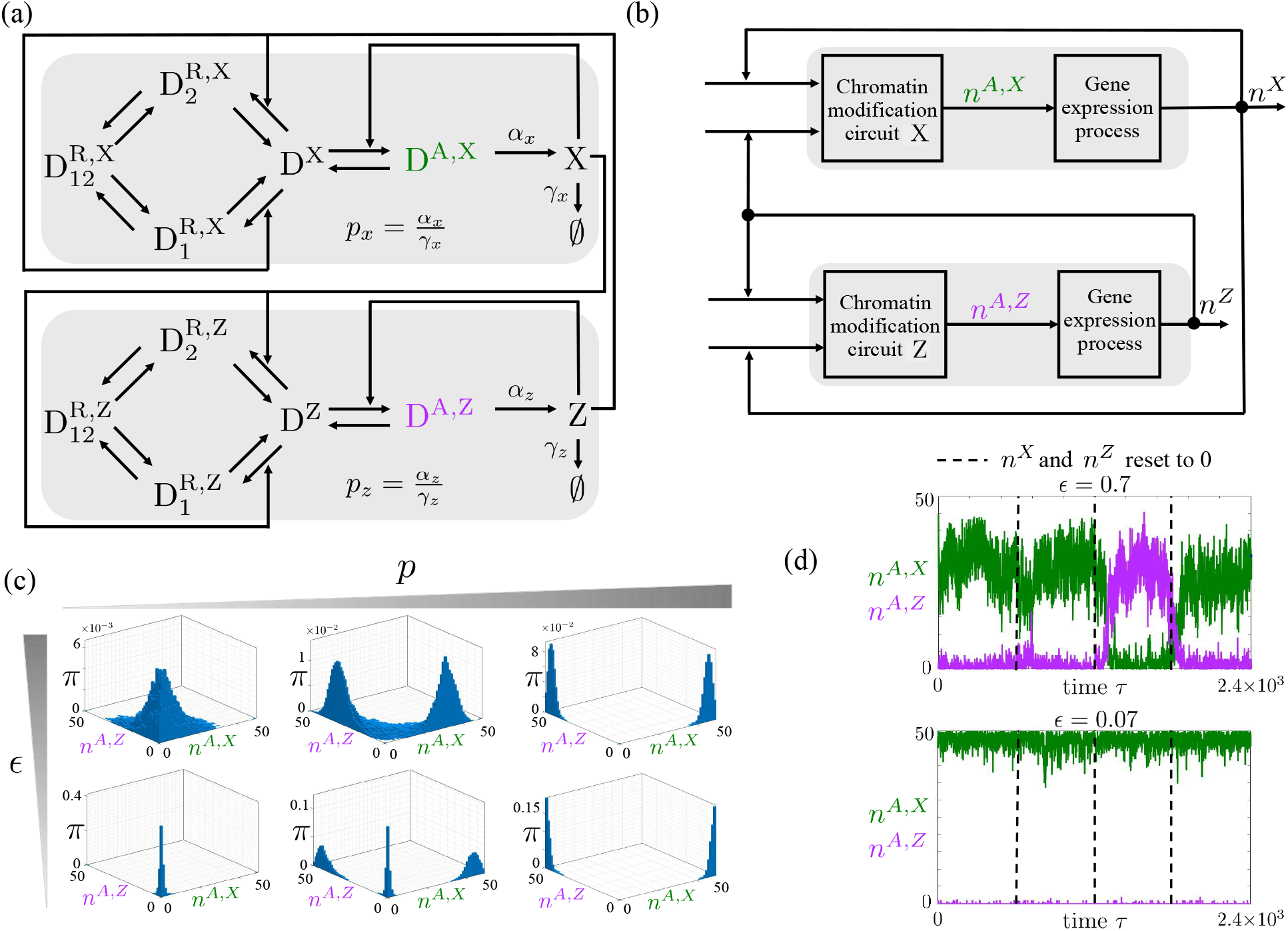
Mutual repression circuit: robust memory of multiple co-existing gene expression patterns. (a) Interaction diagram of two mutually repressing and positively autoregulated genes, wherein the product of each gene recruits writers of repressive chromatin modifications to the other gene. Here, X and Z represent the products of the two genes. (b) Block diagram corresponding to the circuit in panel (a). Here, *n*^*X*^ and *n*^*Z*^ correspond to the number of molecules of X and Z, respectively. (c) Stationary probability distribution *π* of the system obtained by simulating the reactions listed in SI-Tables 8-9 with the SSA, in which *n*^*A,ℓ*^ with *ℓ* = *X, Z* represents the number of nucleosomes in each gene with activating histone modifications. In (d), *p*_*x*_ = *p*_*z*_ = *p* with *p* = 0, 0.1, 10 and *ϵ* = 0.48, 0.2. The parameter values of each plot are listed in SI-Tables 8-9. For all simulations we have *µ* = 1, *µ*′ = 0.6, and *ϵ*′ = 1 (SI-Figures S.32-S.34 show different parameter values). (d) Time trajectories of *n*^*A,X*^ and *n*^*A,Z*^ starting from *n*^*A,X*^ = D_tot_ and *n*^*A,Z*^ = 0, in which *n*^*X*^ and *n*^*Z*^ are reset to zero at the indicated times (dashed line). Time is normalized with respect to 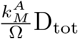. The parameter values for each panel are listed in SI-Tables 10-11. In particular, *p* = 0.15, *µ*′ = 0.6, *µ* = 1, *ϵ*′ = 1 and *ϵ* as indicated. In all plots, we assume equal parameters for both chromatin modification circuits. In our model, *ϵ*, defined in equation (1), is a non-dimensional parameter that quantifies the time scales of basal erasure rate of all modifications relative to those of auto and cross-catalysis. In all plots (*n*^*A,ℓ*^, *n*^*R,ℓ*^) ≈ (50, 0) and (*n*^*A,ℓ*^, *n*^*R,ℓ*^) ≈ (0, 50) correspond to the active and repressed state of gene *ℓ*, with *ℓ* = *X, Z*. In the figure, we use green and purple, respectively, to indicate D^A,X^ and D^A,Z^ and related quantities.

For each gene, the chromatin modification circuit is described by the reactions in Figure 2 and the gene expression reactions are listed in (7). In each gene’s chromatin modification circuit, 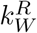 is a monotonically increasing function of the other gene’s product level and 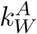 is a monotonically increasing function of the gene’s self product level (SI-Section S.6.1, SI-equations (S.192)). The system is then obtained by input/output connecting two chromatin modification circuits through the TFs that they are expressing (Figure 6(b)).

#### Deterministic behavior

By combining the ODEs of the chromatin modification circuit and those of the gene expression circuit (equations (2) and (8), respectively) for each gene, by defining *X* := *n*^*X*^*/*D_tot_, *Z* := *n*^*Z*^*/*D_tot_ and by properly setting the inputs according to SI-equations (S.195), we obtain the ODEs of the mutual repression system (SI-equations (S.196)). To determine the number and stability of equilibria, we exploit the results obtained for the positive autoregulation circuit. In fact, the block diagram in Figure 6(b) makes it explicit that the mutual repression circuit is the input/output composition of two positively autoregulated genes, in which the output of one gene, *n*^*X*^ or *n*^*Z*^, is used as an input to the other gene by increasing its 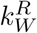. This analysis, described in SI-Section S.6.3, shows that, for low *p* values and large *ϵ*, the system has a unique stable steady state about the origin, where both genes are “off”. By increasing *p*, the system acquires two stable steady states, in which one gene is “on” and the other is “off”. That is, one equilibrium with 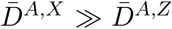 and the other equilibrium with 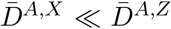. When *ϵ* is small, also the steady state about the origin is stable, while when *ϵ* is large, it is not. Thus, when *p* is large (high expression rate) and *ϵ* is small, the system is tri-stable. When *µ*′ is increased, the system can have four co-existing stable steady states, acquiring a new stable steady state in which both genes are “on” (SI-Figure S.31). Since we expect *µ*′ to be much smaller than one, this last scenario may not be observed in practice.

#### Stochastic behavior

Here, denoting the number of nucleosomes in each gene with activating histone modifications as *n*^*A,X*^ and *n*^*A,Z*^, we first determine how *p*_*x*_, *p*_*z*_, and *ϵ* shape the stationary distribution of (*n*^*A,X*^, *n*^*A,Z*^) and hence the admissible gene expression patterns. Specifically, we perform a computational study of the distribution by simulating the full set of reactions (SI-Table 5) with the SSA [49]. If the production rate constant *p* of both proteins is low, independent of *ϵ*, the distribution has one peak only in correspondence to both genes being “off”, consistent with the deterministic analysis. Therefore, the only possible gene expression pattern in this regime is both genes “off”. When *p* is sufficiently increased, the peak in correspondence of both genes “off” disappears and two peaks arise in correspondence of one gene “on” and the other “off”, thus enabling two symmetric gene expression patterns. There is an intermediate protein production rate constant regime, in which a third gene expression pattern with both genes “off” is also possible. In all cases, decreased *ϵ* makes the distribution more concentrated about each of the peaks, thereby also reducing the probability of transitioning out of that expression pattern (Figure 6(c)). The admissible gene expression patterns are largely shaped by the choice of the two protein production rate constants (SI-Figure S.34). Reducing *µ*′, as before, biases the distribution towards the repressed chromatin state (*n*^*A,X*^, *n*^*A,Z*^) ≈ (0, 0) (SI-Figure S.33) while varying *ϵ*′ does not change the trends with which the key parameters affect the distribution (SI-Figure S.32). The computational results of Figure 6(c) are in agreement with those obtained from the analytical study (SI-Section S.6.4).

Furthermore, we determine computationally how *ϵ* affects the robustness of the memory of an admissible gene expression pattern with respect to resetting the amount of the gene’s protein products X and Z to 0 (Figure 6(d)). Lower *ϵ* values result into enhanced robustness of the pattern’s memory to resetting *n*^*X*^ and *n*^*Z*^ to zero, indicating that the chromatin modification circuit with lower *ϵ* helps memory of co-existing gene expression patterns persist through changes in the availability of TFs to the promoters. Finally, we analyzed whether overexpression of TF can transition the system from one gene expression pattern to the other one. Indeed, even when a gene expression pattern displays high robustness to TFs variability (low *ϵ*), the system allows a transition from one admissible expression pattern to the other one by TF overexpression (SI-Figure S.35). The speed of transition increases with the level of overexpression and decreases with *ϵ*.

## 4 Discussion

We analyzed the dynamics of a biologically motivated circuit motif among histone modifications and DNA methylation within a gene, which we called the gene’s inner chromatin modification circuit (Figure 1(d)). We found that separation of time scales among three constituent processes, basal erasure, recruited erasure, and auto/cross-catalysis controls memory of input stimuli. Specifically, when basal erasure rate *ϵ* is sufficiently lower than both recruited erasure rate *ϵ*′ and auto/cross-catalysis rates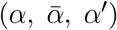, then the chromatin modification circuit shows bistability, hysteresis, and the probability distribution of chromatin state is highly concentrated about the active and repressed states (Figure 3(c)-(d) and Figure 4(a)). Furthermore, the time to memory loss of either active or repressed states increases with *ϵ*, but significantly more so for the repressed chromatin state due to the cross-catalysis among repressive histone modifications and DNA methylation. This asymmetry is enhanced by smaller *µ*′, which is the ratio between the decay rate of DNA methylation and that of activating histone modifications. Since the decay rate of DNA methylation is very small due to the maintenance activity of DNMT1 enzyme, we estimated *µ*′ ≪ 1 (SI-Section S.3.1). In this parameter regime, the active chromatin state becomes poorly robust to repressive input stimuli (Figure 3(f)) and the height of the peak of the distribution corresponding to the active chromatin state decreases (Figure 4(a)). TF-enabled positive autoregulation re-balances this asymmetry, thereby allowing enhanced robustness of the active chromatin state to repressive input stimuli (Figure 5(c)). At the same time, under time scale separation, the chromatin modification circuit enables robustness of the active chromatin state to repetitive disruptions of the positive autoregulation loop (Figure 5(f)). By wiring positively autoregulated chromatin modification circuits, we can thus obtain concurrent mutually exclusive gene expression patterns, which are robust to repeated disruptions of the regulatory links (Figure 6).

In addition to TF-enabled positive autoregulation, other forms of regulation are also possible, such as negative autoregulation and combinations of the two. For example, a form of TF-enabled negative autoregulation is obtained when the protein X recruits writers of repressive chromatin modifications. An instance of a TF that does so is the LANA protein, which recruits DNMT3 to DNA [60]. Furthermore, different forms of autoregulation motifs can be obtained when X regulates the expression of chromatin modifications’ erasers instead of writers. This is observed, for example, for TF Oct4, which increases the expression of H3K9me3 eraser JMJD2A and the expression of TET enzymes [61, 62] or in the regulatory system of olfactory receptor activation, in which the expressed OR protein induces the expression of enzyme Adcy3, which removes the histone demethylase LSD1 [63]. Future work will be devoted to analyzing the dynamical properties of these biologically found regulatory motifs.

The non-dimensional parameters *ϵ, ϵ*′, *µ*, and *µ*′, which control the relative time scales among the circuit’s constituent processes, are known functions of the concentrations of erasers, writers, and readers of nucleosome modifications as well as of cell proliferation rate. As a consequence, our model can predict how experimental interventions on these molecular players and cell division rate will affect plasticity, captured by less concentrated distributions of chromatin states, the kinetics of gene reactivation, and the robustness of active states to repressive stimuli. Indeed, our model predicts that in the absence of time scale separation, which can be obtained by increasing *ϵ*, the cell should be in a more plastic state, and gene reactivation should be faster and less variable. This is consistent with the experiments of [51], in which *ϵ* was artificially increased by increasing proliferation rate. Similarly, our model predicts that increasing *µ*′ should make gene re-activation faster and less stochastic. Our expression 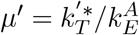 also suggests that increasing the level of TET enzymes effectively increases *µ*′ only if the level of MBD proteins is sufficiently low (SI-equation (S.76)). Indeed, reprogramming experiments where TET was increased did not show dramatic changes on reprogramming kinetics [64], but knocking down MBD proteins showed fast and almost deterministic reactivation kinetics [52]. Future experiments on semi-synthetic chromosomally integrated reporter systems will be required to directly validate some of the model’s predictions. For example, experiments in which *ϵ* is artificially increased by concurrently recruiting erasers for both H3K9 and H3K4 methylation, such as JMJD2A and JARID, will demonstrate the morphing of the distribution from bimodal to unimodal. Experiments where only erasers for H3K4 methylation (JARID) will be recruited can validate the prediction that the memory of the repressed state is extended through the increase of *µ* and *µ*′. Finally, experiments in which H3K9 methylation is completely erased through recruitment of JMJD2A but DNA methylation is present should show loss of bimodality, since this is not present in a model that only includes DNA methylation and H3K4 methylation (SI-Figure S.2). In addition, we will revise the model in order to remove some of the biological simplifications, such as the well-mixed model simplification and the assumption that a nucleosome cannot be characterized by more than one modified histone simultaneously.

Our results suggest that reduced time scale separation (larger *ϵ* and/or reduced *ϵ*′) may be implicated in phenotypic plasticity, wherein larger *ϵ* allows for more probable transitions among multiple admissible gene expression patterns and thus among different phenotypes (Figure 6(c)). While small *ϵ* embodies a gene expression pattern with high robustness to loss of TF binding, larger *ϵ* allows for more probable transitions following TF binding disruptions (Figure 6(d)). It is plausible that physiological requirements for high degree of cellular plasticity at the initial stages of cell fate specification may be satisfied by a transiently larger *ϵ* at cell fate specific genes. Locking-in of the fate may then be aided by a decreased *ϵ* at these cell fate specific genes in terminally differentiated cells [8]. On the other hand, accidental changes of phenotype often result in disease such as cancer [65],[66]. These accidents may correlate with increased *ϵ* and/or *µ*′ at those genes that become disregulated. Global increases in *ϵ* may then be caused by increased proliferation rates since these increase the basal decay rate of all modifications. Indeed, cellular states with high proliferation rates are associated with larger *ϵ* and hence with increased plasticity. These states include fastly proliferating cancer cells, pluripotent stem cells, and, most generally, progenitor cells.

Finally, genomic integration of engineered genetic systems has appeared as a promising approach for the creation of organoids [67] and for *in vivo* transdifferentiation [56, 57]. In particular, we found that positive autoregulation, by recruitment of writers of activating modifications or of erasers of repressive modifications, enables robustness of active chromatin states to endogenous silencing factors. This provides design guidelines for chromosomally integrated gene expression cassettes that are resilient to silencing. Similarly, mutual repression circuits obtained by chromatin state regulation, as opposed to traditional TF-enabled regulation, can be used for future cell fate decision circuits that remain locked-in with high probability but yet are reconfigurable by external interventions.

## Acknowledgements

S.B. was supported by NIH/NIBIB Grant Number R01EB024591 (PI: D.D.V.), in part by NSF Collaborative Research grant MCB-2027949 (PI: D.D.V.) and by the DoD Newton Award for Transformative Ideas during the COVID-19 Pandemic. R.J.W. was supported in part by NSF Collaborative Research grant MCB-2027947 (PI: R.J.W.) and by the Charles Lee Powell Foundation. R.J.W. also gratefully acknowledges the hospitality of the Center of Mathematical Sciences and Applications at Harvard University for her sabbatical visit, during which preliminary discussions were conducted with S.B. and D.D.V. on the research reported here.

## Author contributions

D.D.V. designed the research. S.B. developed the mathematical models. S.B. and D.D.V. performed mathematical analyses. R.J.W. assisted with the mathematical derivation of the stationary probability distribution and time to memory loss expressions. S.B. performed computational simulations. S.B. and D.D.V. analyzed the results. S.B. and D.D.V. wrote the paper.

## References

[1] C. H. Waddington. “The epigenotype”. In: Endeavour (1942).

[2] M. Ptashne. “Epigenetics: Core misconcept”. In: Proc. Natl. Acad. Sci 110 (2013).

[3] Wen Xiong and James E. Ferrell Jr. “A positive-feedback-based bistable ‘memory module’ that governs a cell fate decision”. In: Nature 426 (2003).

[4] N. Carey. The epigenetic revolution. Columbia University Press, 2013.

[5] S. Huang, M. Litt, and C. A. Blakey. Epigenetic Gene Expression and Regulation. Academic Press, 2015.

[6] R. Lanza and A. Atala. Essentials of stem cell biology. Elsevier, 2014.

[7] S. Huang. “Reprogramming cell fates: reconciling rarity with robustness”. In: BioEssays 31 (2009).

[8] J. Holmberg and T. Perlmann. “Maintaining differentiated cellular identity”. In: Nature Reviews 13 (2012).

[9] Sui Huang et al. “Bifurcation dynamics in lineage-commitment in bipotent progenitor cells”. In: Developmental Biology 305.2 (2007), pp. 695–713. ISSN: 0012-1606. doi: http://dx.doi.org/10.1016/j.ydbio.2007.02.036. URL: http://www.sciencedirect.com/science/article/pii/S0012160607001674.

[10] C. D. Allis et al. Epigenetics. Cold Spring Harbor Laboratory Press, Second Edition, 2015.

[11] Edith Heard and Robert A. Martienssen. “Transgenerational Epigenetic Inheritance: Myths and Mechanisms”. In: Cell 157 (2014).

[12] C. D. Allis and T. Jenuwein. “The molecular hallmarks of epigenetic control”. In: Nature Reviews 17 (2016).

[13] R. Holliday and J. E. Pugh. “DNA Modification Mechanisms and Gene Activity during Development”. In: American Association for the Advancement of Science 187 (1975).

[14] A D Riggs. “X inactivation, differentiation, and DNA methylation”. In: Cytogenet Cell Genet. 14 (1975).

[15] J. Du et al. “DNA methylation pathways and their crosstalk with histone methylation”. In: Nat Rev Mol Cell Biol 16 (2015).

[16] L. Ordovas et al. “Efficient recombinase-mediated cassette exchange in hpscs to study the hepatocyte lineage reveals aavs1 locus-mediated transgene inhibition”. In: Stem Cell Reports 5 (2015).

[17] S. Y. Alhaji, S. C. Ngai, and S. Abdullah. “Silencing of transgene expression in mammalian cells by DNA methylation and histone modifications in gene therapy perspective”. In: Biotechnology and Genetic Engineering Reviews 35 (2019).

[18] M. Fitzgerald et al. “Rosa26 docking sites for investigating genetic circuit silencing in stem cells”. In: Synthetic Biology 5 (2020).

[19] J. Zimak et al. “Epigenetic silencing directs expression heterogeneity of stably integrated multi-transcript unit genetic circuits”. In: Sci Rep 11 (2021).

[20] Leonie Ringrose and Martin Howard. “Dissecting chromatin-mediated gene regulation and epigenetic memory through mathematical modelling”. In: Current Opinion in Systems Biology 3 (2017).

[21] Maxim N. Artyomov, Alexander Meissner, and Arup K. Chakraborty. “A Model for Genetic and Epigenetic Regulatory Networks Identifies Rare Pathways for Transcription Factor Induced Pluripotency”. In: PLOS Comput. Biol. 6 (2010).

[22] Max Flöttmann, Till Scharp, and Edda Klipp. “A stochastic model of epigenetic dynamics in somatic cell reprogramming”. In: Frontiers in Physiology (2012).

[23] S. S. Ashwin and Masaki Sasai. “Effects of Collective Histone State Dynamics on Epigenetic Landscape and Kinetics of Cell Reprogramming”. In: Nature Reports (2015).

[24] T. Chen, M. A. Al-Radhawi, and E. D. Sontag. “A mathematical model exhibiting the effect of DNA methylation on the stability boundary in cell-fate networks”. In: Epigenetics 16.4 (2021), pp. 436–457.

[25] W. Jia et al. “A possible role for epigenetic feedback regulation in the dynamics of the epithelial-mesenchymal transition (EMT)”. In: Phys Biol 16.6 (2019).

[26] Ian B. Dodd et al. “Theoretical Analysis of Epigenetic Cell Memory by Nucleosome Modification”. In: Cell 129 (2007).

[27] Tianyi Zhang, Sarah Cooper, and Neil Brockdorff. “The interplay of histone modifications – writers that read”. In: EMBO Reports 16 (2015).

[28] Joanna Wysocka et al. “Human Sin3 deacetylase and trithorax-related Set1/Ash2 histone H3-K4 methyltransferase are tethered together selectively by the cell-proliferation factor HCF-1”. In: GENES & DEVELOPMENT 17 (2003).

[29] Olivier Binda et al. “Trimethylation of histone H3 lysine 4 impairs methylation of histone H3 lysine 9”. In: Epigenetics 5 (2010).

[30] B. Jin, Y. Li, and K. D. Robertson. “DNA Methylation. Superior or Subordinate in the Epigenetic Hierarchy?” In: Genes Cancer 2.5 (2011), pp. 607–617.

[31] Steen K. T. Ooi et al. “DNMT3L connects unmethylated lysine 4 of histone H3 to de novo methylation of DNA”. In: Nature 448 (2007).

[32] A. J. Courey. Mechanisms in Trancriptional Regulation. Blackwell Publishing, 2008.

[33] TaeSoo Kim and Stephen Buratowski. “Dimethylation of H3K4 by Set1 Recruits the Set3 Histone Deacetylase Complex to 50 Transcribed Regions”. In: Cell 137 (2009).

[34] Yen-Sin Ang et al. “Wdr5 Mediates Self-Renewal and Reprogramming via the Embryonic Stem Cell Core Transcriptional Network”. In: Cell 145 (2011).

[35] Tomas Stopka et al. “PU.1 inhibits the erythroid program by binding to GATA-1 on DNA and creating a repressive chromatin structure”. In: The EMBO Journal 24 (2005).

[36] Nathaniel A. Hathaway et al. “Dynamics and Memory of Heterochromatin in Living Cells”. In: Cell 149 (2012).

[37] M. Trerotola et al. “Epigenetic inheritance and the missing heritability”. In: Human Genomics 9 (2015).

[38] C. Alabert et al. “Two distinct modes for propagation of histone PTMs across the cell cycle”. In: Research Communication, CSHL (2015).

[39] Anup K. Upadhyay et al. “Coordinated Methyl-lysine Erasure: Structural and Functional Linkage of a Jumonji demethylasedomain and a Reader domain”. In: Curr Opin Struct Biol 21 (2011).

[40] Chunlei Jin et al. “TET1 is a maintenance DNA demethylase that prevents methylation spreading in differentiated cells”. In: Nucleic Acids Research 42 (2014).

[41] Kasper Dindler Rasmussen and Kristian Helin. “Role of TET enzymes in DNA methylation, development, and cancer”. In: GENES & DEVELOPMENT 30 (2016).

[42] Anne K. Ludwig et al. “Binding of MBD proteins to DNA blocks Tet1 function thereby modulating transcriptional noise”. In: Nucleic Acid Research 45 (2017).

[43] Bergstrom CT Genereux DP Miner BE and Laird CD. “A population-epigenetic model to infer site-specific methylation rates from double-stranded DNA methylation patterns”. In: Proc. Natl. Acad. Sci 102 (2005).

[44] L Lopez-Serra and M Esteller. “Proteins that bind methylated DNA and human cancer: reading the wrong words”. In: British Journal of Cancer 98 (2008).

[45] X. et al. Nan. “Transcriptional repression by the methyl-CpG-binding protein MeCP2 involves a histone deacetylase complex”. In: Nature 393 (1998).

[46] P.L. et al. Jones. “Methylated DNA and MeCP2 recruit histone deacetylase to repress transcription”. In: Nature Genetics 19 (1998).

[47] Francois Fuks et al. “The Methyl-CpG-binding Protein MeCP2 Links DNA Methylation to Histone Methylation”. In: The J. of Biological Chemistry 278 (2003).

[48] FrancEöis Fuks et al. “The DNA methyltransferases associate with HP1 and the SUV39H1 histone methyltransferase”. In: Nucleic Acids Research 31 (2003).

[49] Daniel T. Gillespie. “Stochastic Simulation of Chemical Kinetics”. In: Annual Review of Physical Chemistry 58.1 (2007). PMID: 17037977, pp. 35–55. doi: 10.1146/annurev.physchem.58.032806.104637.

[50] Lacramioara Bintu et al. “Dynamics of epigenetic regulation at the single-cell level”. In: Science (2016).

[51] Jacob Hanna et al. “Direct cell reprogramming is a stochastic process amenable to acceleration”. In: Nature 462 (2009).

[52] Yoach Rais1 et al. “Deterministic direct reprogramming of somatic cells to pluripotency”. In: Nature (2013).

[53] Liling Tang, Eva Nogales, and Claudio Ciferri. “Structure and Function of SWI/SNF Chromatin Remodeling Complexes and Mechanistic Implications for Transcription”. In: Prog Biophys Mol Biol. 102 (2010).

[54] L. A. Boyer et al. “Core Transcriptional Regulatory Circuitry in Human Embryonic Stem Cells”. In: Cell 122 (2005), pp. 947–956.

[55] Stewart H. Lecker, Alfred L. Goldberg, and William E. Mitch. “Protein Degradation by the Ubiquitin–Proteasome Pathway in Normal and Disease States”. In: Journal of the American Society of Nephrology 17 (2006).

[56] C. Jopling, S. Boue, and J. C. I. Belmonte. “Dedifferentiation, transdifferentiation and reprogramming: three routes to regeneration”. In: Nature Reviews Molecular Cell Biology 12 (2011).

[57] A. Cie,,slar-Pobuda et al. “Transdifferentiation and reprogramming: Overview of the processes, their similarities and differences”. In: Biochimica et Biophysica Acta (BBA) - Molecular Cell Research 1864 (2017).

[58] Pavel Burda et al. “GATA-1 Inhibits PU.1 Gene via DNA and Histone H3K9 Methylation of Its Distal Enhancer in Erythroleukemia”. In: PLOS ONE (2015).

[59] A. Ralston and J. Rossant. “Genetic regulation of stem cell origins in the mouse embryo”. In: Clinical Genetics 68.2 (2005), pp. 106–112. doi: 10.1111/j.1399-0004.2005.00478.x.

[60] M. Shamay et al. “Recruitment of the de novo DNA methyltransferase Dnmt3a by Kaposi’s sarcoma-associated herpesvirus LANA”. In: Proceedings of the National Academy of Sciences 103.39 (2006), pp. 14554–14559.

[61] Yuin-Han Loh et al. “Jmjd1a and Jmjd2c histone H3 Lys 9 demethylases regulate self-renewal in embryonic stem cells”. In: GENES & DEVELOPMENT 21 (2007).

[62] Y. Wu, Z. Guo, and Y. Liu et al. “Oct4 and the small molecule inhibitor, SC1, regulates Tet2 expression in mouse embryonic stem cells”. In: Mol Biol Rep 40 (2013).

[63] X.J. Tian et al. “Achieving diverse and monoallelic olfactory receptor selection through dual-objective optimization design”. In: Proc Natl Acad Sci U S A 113.21 (2016), E2889–E2898.

[64] Y. Gao et al. “Replacement of Oct4 by Tet1 during iPSC Induction Reveals an Important Role of DNA Methylation and Hydroxymethylation in Reprogramming”. In: Cell Stem Cell (2013).

[65] J. M. W. Slack. “Metaplasia and transdifferentiation: from pure biology to the clinic”. In: Nature Rev. Mol. Cell Bio. 8 (2007).

[66] J. Holmberg et al. “Activation of neural and pluripotent stem cell signatures correlates with increased malignancy in human glioma”. In: PLoS ONE 6 (2011).

[67] S.M. Chuva de Sousa Lopes et al. E. Garreta R.D. Kamm. “Rethinking organoid technology through bioengineering”. In: Nat. Mater. 20 (2021).

[68] D. Del Vecchio and R. M. Murray. Biomolecular Feedback Systems. Princeton University Press, 2014.

